# eRNA profiling uncovers the enhancer landscape of oesophageal adenocarcinoma and reveals new deregulated pathways

**DOI:** 10.1101/2022.05.11.491502

**Authors:** Ibrahim Ahmed, Shen-Hsi Yang, Samuel Ogden, Wei Zhang, Yaoyong Li, the OCCAMS consortium, Andrew D. Sharrocks

## Abstract

Cancer is driven by both genetic and epigenetic changes that impact on gene expression profiles and the resulting tumourigenic phenotype. Enhancers are transcriptional regulatory elements that are key to our understanding of how this rewiring of gene expression is achieved in cancer cells. Here we have harnessed the power of RNA-seq data from hundreds of patients with oesophageal adenocarcinoma (OAC) or its precursor state Barrett’s oesophagus (BO) coupled with open chromatin maps to identify potential enhancer RNAs (eRNAs) and their associated enhancer regions in this cancer. We identify ∼1000 OAC-specific enhancers and use this data to uncover new cellular pathways that are operational in OAC. Among these are enhancers for *JUP, MYBL2* and *CCNE1*, and we show that their activity is required for cancer cell viability. We also demonstrate the clinical utility of our dataset for identifying disease stage and patient prognosis. Our data therefore identify an important set of regulatory elements that enhance our molecular understanding of OAC and point to potential new therapeutic directions.

## Introduction

Enhancers are distal regulatory elements that generally promote gene expression by engaging with the promoters of their target genes in *cis* (Andersson and Sandelin, 2020), although they have also been observed acting in *trans* (Hu et al, 2008; Spilianakis et al, 2005) and more recently, through hubs of activity on extra-chromosomal DNA species (Hung et al, 2021). Active enhancers are characterised by the presence of histone marks such as H3K27ac and H3K4me1 (Creyghton et al, 2010; Heintzman et al, 2007). However, it has been shown that enhancers can be the site of production for small transcripts termed enhancer RNAs (eRNAs) (De Santa et al, 2010; Kim et al, 2010). Whilst the functionality of eRNAs is still under debate, there is a large body of evidence associating the production of eRNAs with enhancer activity, and subsequent target gene activation (Tyssowski et al, 2018; Andersson et al, 2014; Chen et al, 2018). This association with gene expression has allowed eRNA-defined enhancer activity to serve as a specific marker for developmental stage and tissue type (Yan et al, 2019; Huang et al, 2016). Furthermore, eRNAs provide molecular markers for disease, often with more sensitivity than the gene expression pattern itself (Zhang et al, 2019; Chen et al, 2018).

During tumourigenesis, there are widespread changes to gene expression patterns that are associated with rewiring of the regulatory landscape in an enhancer-driven manner (Li et al, 2015; Hsieh et al, 2014). This can be accompanied by eRNA production. For example, production of an eRNA from the *PSA* gene enhancer is associated with increased *PSA* expression in castration-resistant prostate cancer (Zhao et al, 2016). This potentially makes the production of eRNAs a biomarker for cancer, but this is widely underappreciated. Indeed, a recent pan cancer study demonstrated that eRNAs can serve as prognostic markers across various cancer types and provide novel insights into cancer biology (Chen et al, 2018), leading to the identification of therapeutic opportunities.

Oesophageal adenocarcinoma (OAC) has an overall 5-year survival rate of approximately 15%, making it a leading global cause of cancer-associated deaths (Coleman et al, 2018). OAC is believed to arise in a stepwise fashion from the pre-cancerous lesion Barrett’s oesophagus (BO) (Peters et al, 2019). A number of large-scale DNA sequencing studies have been performed into the pathogenesis of OAC from Barrett’s (Frankell et al, 2019; Ross-Innes et al, 2015; Stachler et al, 2015), however there is still uncertainty concerning the precise molecular mechanisms. In an effort to understand potential epigenetic contributors to OAC, we have previously demonstrated that changes in chromatin accessibility play a role in the transition to OAC (Britton et al, 2017; Rogerson et al, 2019; Rogerson et al, 2020). These chromatin changes are often associated with non-coding regions of the genome that may represent regulatory elements such as enhancers.

Here, we build on our previous work identifying chromatin changes during the BO to OAC transition (Britton et al, 2017; Rogerson et al, 2019; Rogerson et al, 2020). By integrating total RNA-seq data from BO and OAC patients generated by the Oesophageal Cancer Clinical and Molecular Stratification (OCCAMS) consortium dataset, with previously generated chromatin accessibility data on these tissues (Britton et al., 2017; Rogerson et al., 2019; Rogerson et al., 2020; Corces et al., 2018), we identify, and validate, pervasive eRNA production at regions of accessible chromatin, indicative of enhancer activity. Subsequent interrogation of genes associated with these enhancers identified deregulated pathways of importance and demonstrated the potential clinical utility of eRNA profiling in OAC.

## Results

### Identification of potential intergenic eRNA transcripts in OAC patients

eRNAs are generally unstable and lowly abundant, making them hard to detect in RNAseq data sets. We therefore harnessed the sequencing power derived from combining hundreds of patient samples to discover potential eRNAs (Fig. 1A). To identify eRNAs that are relevant to OAC we interrogated RNA-seq data generated from 210 OAC patients and also from 108 Barrett’s patients to identify differentially upregulated eRNAs in OAC (Jammula et al., 2020). After mapping sequencing reads to the genome, we excluded all regions corresponding to gene bodies as well as sequences 2 kb upstream from the transcriptional start site (TSS) and 500 bp downstream from the annotated transcriptional termination site (TTS) (Supplementary Fig. 1A) to avoid interference from promoter sequences and read through transcription respectively. Next, we identified all of the accessible regions of chromatin in OAC and Barrett’s samples within this truncated genome by creating a union peak set from ATAC-seq performed on 14 OAC and 4 Barrett’s samples (Britton et al., 2017; Rogerson et al., 2019; Rogerson et al., 2020; Corces et al., 2018) (resulting in 150,265 peaks). To focus on potential enhancer regions, we then took the RNA-seq data from both classes of patients and assessed raw reads wihtin these accessible chromatin regions. This identified 61,349 intergenic regions that contain RNA transcripts and represent potential eRNA regions. We then filtered these based on a read count ≥ 3 and FPM value ≥ 1.5, which resulted in a final high confidence set of 4600 potential eRNA containing enhancer regions in OAC and Barrett’s patients (Supplementary Table S1).

**Figure 1.**
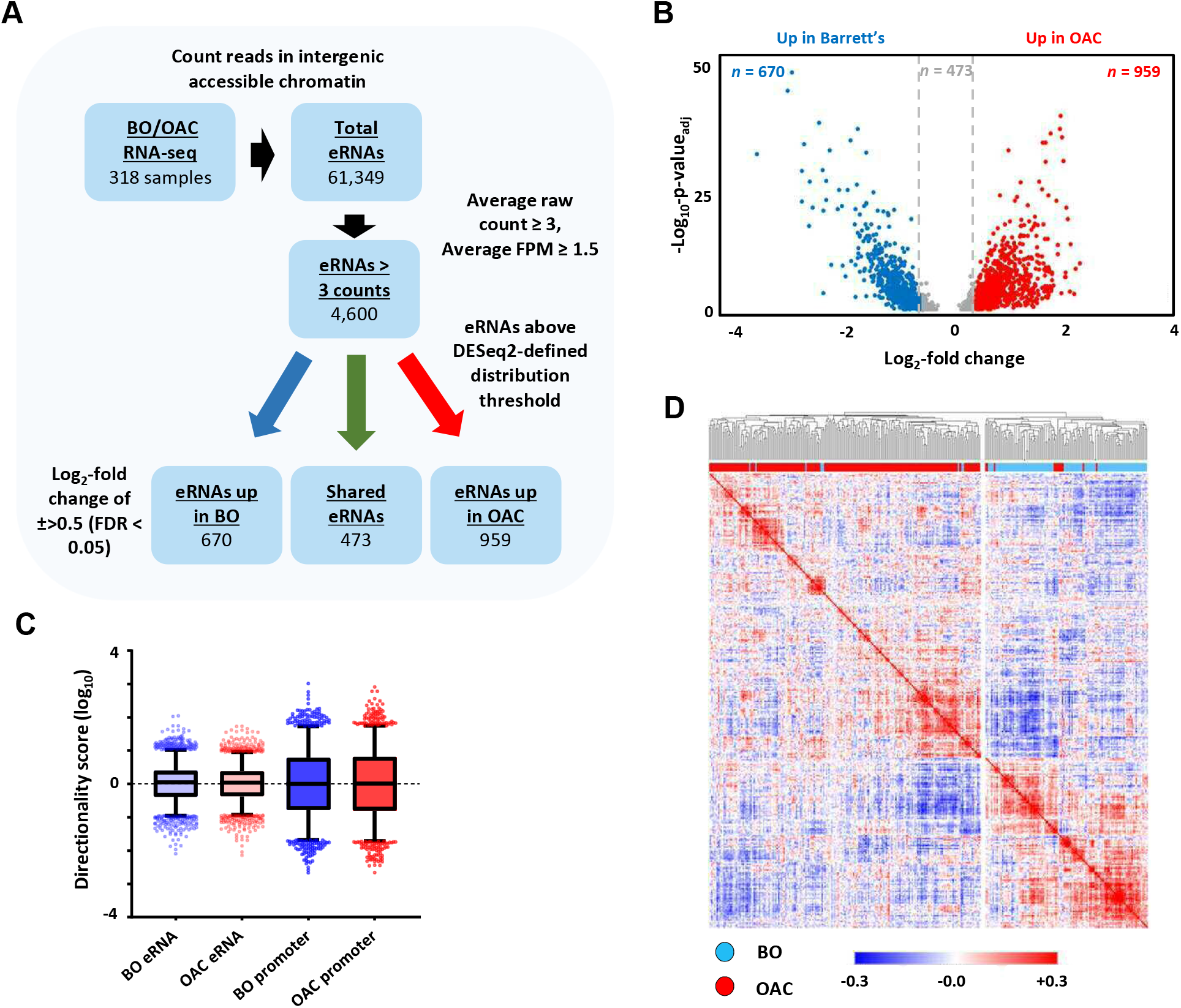
Identification of enhancer transcription in OAC and Barrett’s patients. (A) eRNA identification strategy. The numbers of putative eRNAs identified at each stage are indicated. (B) Volcano plot displaying the differentially expressed (± Log_2_FC 0.5, < *p*_adj_ = 0.05) eRNAs (*n* = 2,102). (C) Directionality scores for BO-or OAC-specific eRNAs compared to promoters (D) Pearson’s correlation and hierarchical clustering of BO *(n* = 108) and OAC *(n* = 210) patient tissue total RNA-seq samples according to row z-score normalised expression levels in the 4600 eRNA regions.

Next, we identified differentially transcribed eRNAs in each disease state (+/->0.5 log_2_ fold; P-value_adj_ <0.05; Fig. 1B; Supplementary Fig. S1A) and found 959 to be significantly upregulated in OAC (Supplementary Table S2) and 670 to be more active in Barrett’s patients (Supplementary Table S3). 2,498 eRNAs did not meet the stringent DESeq2 distribution threshold and were discarded. The remaining putative eRNAs exhibited low directionality score distributions consistent with the bi-directional transcription associated with eRNA production (Fig. 1C; Supplementary Fig. S1B).

To probe the clinical utility of eRNA region profiling, we analysed the expression levels found in the 4,600 eRNA regions across all of the OAC and Barrett’s samples. Hierarchical clustering showed a clear separation of Barrett’s and OAC patients (Fig. 1D). While clustering based on whole RNA-seq data gave broadly similar separation of OAC and Barrett’s samples, several more were misclassified compared to eRNA-based profiling (Supplementary Fig. S1C; 37 versus 8 samples). Clustering of the expression of the same eRNA regions in a different RNA-seq dataset (Maag et al., 2017) also provided a good separation of the OAC and Barrett’s samples (Supplementary Fig. S1D), further demonstrating the relevance of this dataset. In summary therefore, we have identified a panel of potential eRNA generating regions that can be used for discriminating OAC samples from the pre-cancerous Barrett’s state.

### eRNA-associated regions show enhancer-like characteristics

Having identified a group of accessible chromatin regions expressing potential eRNAs, we next sought further evidence to associate these with enhancer-like activity. First, we examined chromatin accessibility in OAC patients and found higher levels at eRNA expressing loci compared to a random set of open chromatin regions (Fig. 2A; left). Furthermore, these regions also exhibited higher levels of the H3K27ac chromatin mark in OE19 OAC-derived cells that is usually associated with active enhancers (Fig. 2A; right). This was not just a function of increased accessibility as a control group of more highly accessible regions did not exhibit increased levels of H3K27ac (Supplementary Fig. S2A). Indeed, chromatin accessibility levels are a weak indicator of eRNA transcription levels across patient samples, as even regions of greater accessibility contained much lower transcript levels (Fig. 2B; Supplementary Fig. S2B). Thus, eRNA regions are more accessible but the reciprocal is not true, as more accessible regions do not necessarily show higher levels of eRNA transcription.

**Figure 2.**
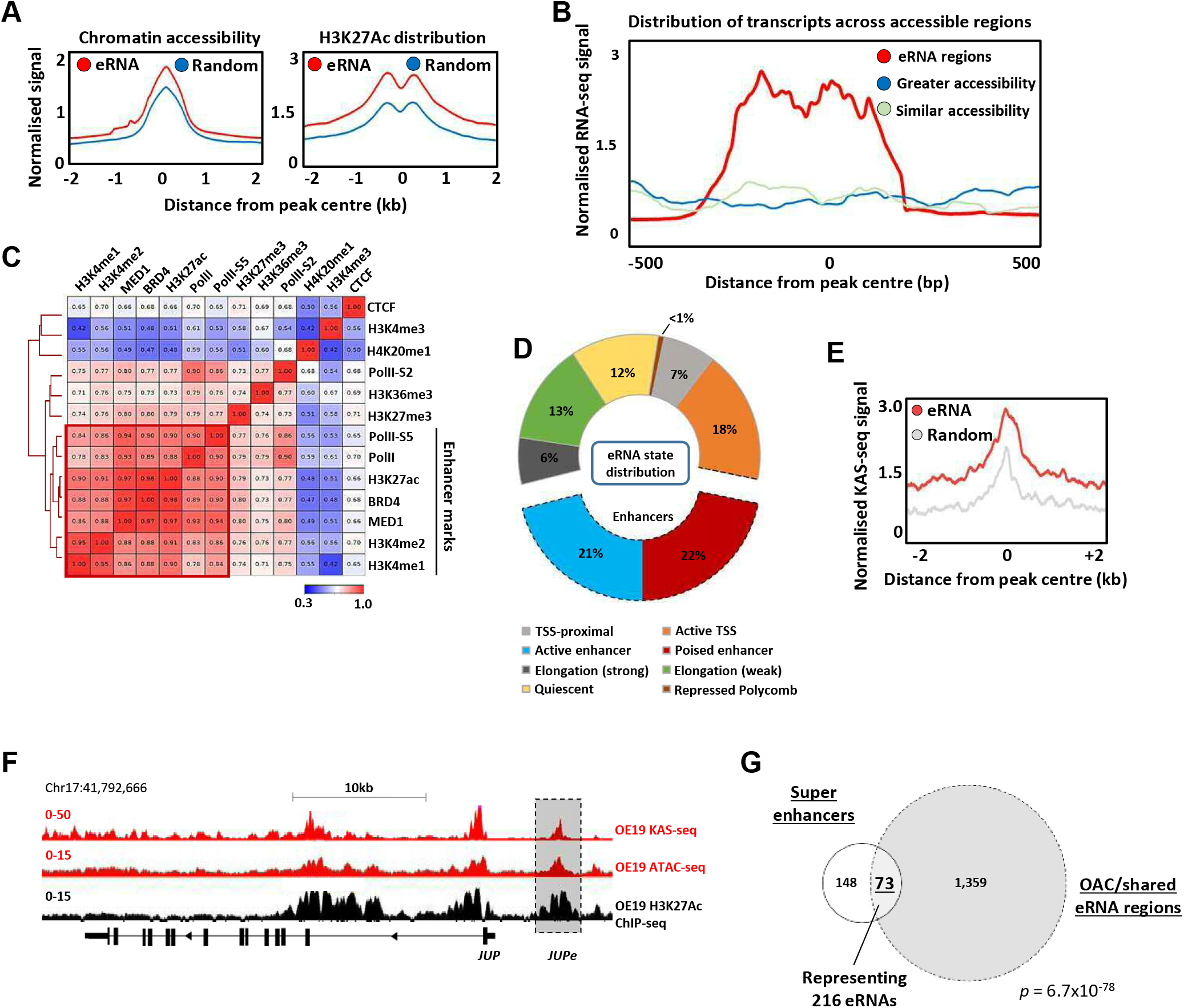
Putative eRNAs are associated with enhancer-like genomic regions. (A) Metaplots of (top left) patient tissue chromatin accessibility and (bottom left) OE19 cell H3K27ac ChIP-seq signal at all 4,600 eRNA regions compared to 4,600 random regions of accessible chromatin. (B) Distribution of transcription at 4,600 eRNA regions compared to 4,600 randomly selected regions of similar or greater chromatin accessibility (regions shown as peak centre ± 0.5 kb). (C) Pearson’s correlation and hierarchical clustering of CUT&TAG signal at 4,600 eRNA regions for various chromatin-associated factors. (D) Distribution of ChromHMM emission states for 4,600 eRNA regions. (E) Metaplots of KAS-seq signal in OE19 cells at 4,600 eRNA regions compared to 4,600 random regions of accessible chromatin. (F) Genome browser view of OE19 KAS-seq, OE19 ATAC-seq data, and OE19 H3K27ac ChIP-seq at the *JUP* locus with the *JUPe* eRNA highlighted. (G) Venn-diagram of overlap between 221 high-confidence intergenic super enhancers and 1,432 eRNAs (specific to OAC or shared with BO eRNA ; *p*-value is shown; hypergeometric test).

To provide further evidence for association with active enhancers in OAC cells, we used CUT&TAG (Kaya-Okur et al., 2020) to profile a range of histone marks and chromatin associated proteins in OE19 cells and correlated the levels of these across the eRNA expressing regions (Fig. 2C). There is clear co-association with a range of enhancer associated marks and proteins, including RNA polymerase II, BRD4 and the MED1 subunit of mediator. This is also evident when visualising the data as heat maps compared to random regions of accessible chromatin, where there are higher levels of the chromatin associated marks/proteins that categorise enhancers in the eRNA regions (Supplementary Fig. S2C). Conversely, there is a clear depletion of the promoter associated mark H3K4me3 (Supplementary Fig. S2D). We also examined the distribution of the chromatin marks H3K4me1 (associated with active and poised enhancers) and H3K4me3 (associated with active promoters) found in gastric adenocarcinoma patients (GAC) which are molecularly similar to OAC patients (The Cancer Genome Atlas, 2017). Neither mark is enriched in the eRNA containing regions compared to random accessible regions (Supplementary Fig. S2E). However, there is a clear enrichment of H3K4me1 and depletion of H3K4me3 in eRNA containing regions compared to promoters, consistent with their designation as potential enhancers (Supplementary Fig. S2F). Conversely, the CpG content of the eRNA regions was substantially lower than at promoter regions (Supplementary Fig. S2G).

To further probe links between eRNA containing regions and enhancer activity, we partitioned the genome into a series of states via a Hidden Markov Model (HMM states) (Supplementary Fig. 2H). Positional information was used to mark these HMM states as promoter proximal or distal (Supplementary Fig. S2I). Re-evaluation of the eRNA expressing regions showed that 43% were associated with regions designated as enhancers (Fig. 2D), with very few regions designated as quiescent or repressed (13% compared to 26% in randomly selected open chromatin regions; Supplementary Fig. S2J).

The co-association of eRNA expressing regions with genomic elements associated with active chromatin marks is strongly suggestive of enhancer-like activity. However, to provide further evidence for active ongoing transcription at these loci we performed KAS-seq (Wu et al., 2020) in OE19 cells to identify areas of DNA strand opening as observed in the transcription bubble. The three replicates showed good congruency (Supplementary Fig. S3A), and we merged all three to call peaks of DNA strand opening (Supplementary Table S4). These peaks show a highly significant overlap with the eRNA regions we identified from patient samples (Supplementary Fig. S3B), and higher levels of KAS-seq signal are associated with eRNA regions compared to random regions (Fig. 2E). This is exemplified by eRNAs associated with a putative enhancer region located upstream from *JUP* (Fig. 2F), *CCNE1* and *MYBL2* gene loci (Supplementary Fig. S3C and S3D). Furthermore, when we combined eRNA regions with regions showing KAS-seq peaks, higher levels of H3K27ac were observed compared to those lacking concomitant KAS-seq signal (Supplementary Fig. S3E), indicative of higher activity.

Finally, we focussed on potential super enhancers as these have been shown to play important roles in cancer-specific gene regulation (Hnisz et al., 2013) including gastroesophageal cancers (Ooi et al., 2016). We identified potential super enhancers in OE19 cells using HOMER (Heinz et al., 2010; Whyte et al., 2013) from peak sets generated from H3K27ac ChIP-seq (for histone activation marks) and ATAC-seq (for open chromatin) (Supplementary Fig. S4A; Supplementary Table S5). We then overlapped these peak sets to generate a dataset with both indicators of super enhancer activity, excluding promoter regions from our analysis. This resulted in 221 high confidence super enhancers (Supplementary Fig. S4B). Next, the constituent enhancers within these super enhancers were overlapped with the eRNA regions we identified from OAC patients, producing a final list of 73 super enhancers showing evidence of eRNA activity, with a total of 216 eRNA regions residing in these super enhancers (Fig. 2G). Multiple eRNA regions identified in patient samples are therefore associated with super enhancers as exemplified by the *ELF3* super enhancer (Supplementary Fig. S4C). The genes associated with these super enhancers are enriched in several biological processes with direct relevance to OAC, such as MAPK signalling and cadherin binding (Supplementary Fig. S4D).

Collectively, these data strongly indicate that the eRNA associated regions we discovered in patient samples represent areas of enhancer activity due to the presence of enhanced accessibility, enhancer associated chromatin marks/proteins, and evidence for actively transcribing RNA polymerase in OAC cells.

### Association of eRNA regions to target genes and regulatory transcription factors

Next, we asked whether we could identify upstream transcription factors that might control eRNA levels and provide insights into the regulatory landscape of OAC. First, we identified binding motifs for transcription factors that are over-represented in OAC or Barrett’s-specific eRNA producing regions. This revealed enrichment for transcription factors previously identified as relevant for OAC including AP1, KLF5 and HNF1 (Britton et al., 2017; Rogerson et al., 2019; Rogerson et al., 2020; Chen et al., 2020) as well as CTCF, a factor implicated in enhancer activity (Ren et al., 2017) (Fig. 3A: Supplementary Table S6A). However, the frequency of motif occurrence differed between eRNA-defined and open chromatin-defined enhancers in OAC patients; HNF1 motifs were significantly more enriched in eRNA-defined enhancers whereas AP1 and KLF5 motifs were more enriched in enhancers defined by increased accessibility alone (Fig. 3B). AP1 motifs were again identified in Barrett’s-specific regions as well as a different set of motifs including p53 binding motifs (Fig. 3A; Supplementary Table S6B). Similarly, calculating differential binding scores revealed higher binding activity of AP1 and HNF1 in OAC-specific regions and conversely higher p53 binding activity in Barrett’s specific regions (Fig. 3C; Supplementary Table S7). Thus, eRNA-defined enhancers reveal the activity of disease stage-specific transcriptional regulators. To further explore this point, we sought evidence for enhancer occupancy by transcription factors in OAC cells and found substantially more binding signal of KLF5 derived from ChIP-seq in OE19 cells (Rogerson et al., 2020) for eRNA-defined regions with KLF5 motifs compared to regions lacking the motif, or control genomic regions (Fig. 3D). Furthermore, evidence for KLF5-mediated regulation was obtained by the significant overlap between the genes associated with the same eRNA regions (ie containing KLF5 motifs) and those genes downregulated upon KLF5 depletion in OE19 cells (Fig. 3E; Supplementary S5A-C). We also examined the effect of AP1 inhibition by expressing a dominant-negative FOS derivative (dnFOS; Olive et al., 1997) in OE19 cells and compared the downregulated gene profile with the eRNA-associated genes containing AP1 motifs that we have identified. Again, we observed a significant overlap between the genes associated with eRNA regions containing AP1 motifs in OAC samples and those genes downregulated upon AP1 inhibition (Fig. 3F; Supplementary Fig. S5D-F). Thus, both motif discovery and functional dissection demonstrates that KLF5 and AP1 are likely major players in eRNA-defined enhancer activation in OAC patients.

**Figure 3.**
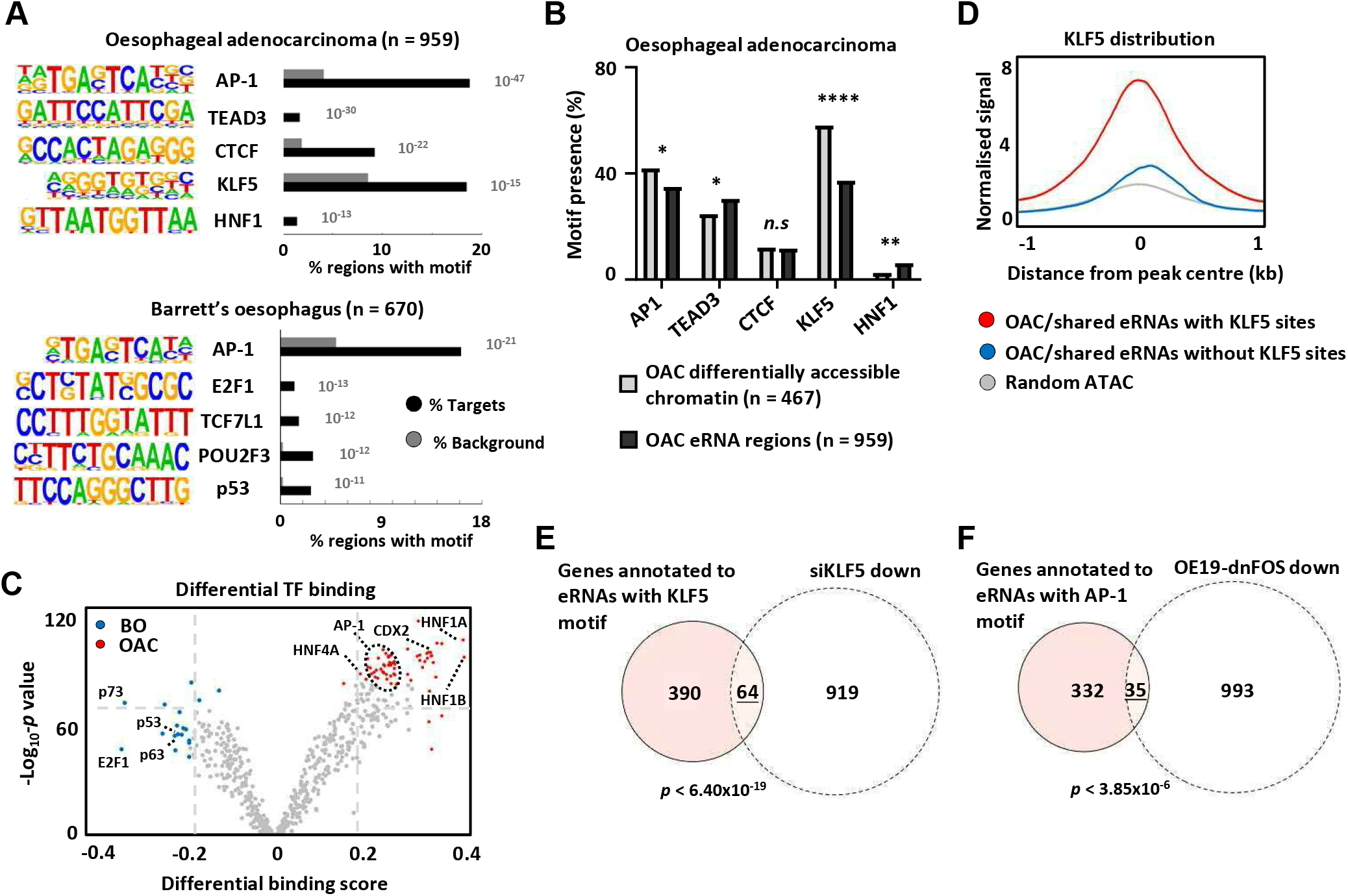
Association of eRNA regions with transcriptional regulators. (A) Transcription factor *de novo* motif enrichment using HOMER, at eRNAs differentially expressed in OAC (top; n = 959) and Barrett’s (bottom; n = 670) (p-values are shown). (B) Bar graphs displaying the frequency of motif prevalence of the top five enriched motifs at eRNA regions differentially expressed in OAC (top; n = 959) compared to differentially accessible intergenic chromatin (**** = *p* < 0.0001; ** = *p* < 0.01; * = *p* < 0.05; N-1 Chi-squared test). (C) Volcano plot showing differential TF binding (± 0.2 differential binding score or 2: -log_10_ padj 70) at 4,600 eRNAs regions using TOBIAS (Bentsen et al., 2020). (D) Metaplots of KLF5 ChIP-seq signal from OE19 cells at eRNAs (specific to OAC or shared with BO eRNA) containing a KLF5 motif, lacking a KLF5 motif or randomly selected open chromatin regions. (E) Venn diagram displaying overlap between genes annotated to KLF5 motif containing eRNAs (specific to OAC or shared with BO eRNA) with genes downregulated upon siKLF5 treatment (Log_2_FC 2:1.0, < padj = 0.05) in OE19 cells (*p-*value is shown; Fisher’s exact test). (F) Venn diagram displaying overlap between genes annotated to AP-1 motif containing eRNAs (specific to OAC or shared with BO eRNA) with genes downregulated upon dnFOS induction (Log_2_FC 2:0.5, < padj = 0.05) in OE19 cells (*p-*value is shown; Fisher’s exact test).

To understand the potential biological consequences of eRNA activation, we then linked the differentially active eRNA regions to putative target genes with the nearest gene model using HOMER (Heinz et al., 2010) resulting in 528 genes in OAC and 380 genes in Barrett’s. Gene ontology (GO) analysis identified several terms relevant to the OAC phenotype such as “Cell migration” and “MAPK signalling” whereas Barrett’s-specific regions identified genes associated with various metabolic processes and “epithelial differentiation” that would be expected for this intestinal metaplastic tissue (Fig. 4A). We also performed differential gene expression analysis on the whole RNA-seq datasets to identify genes preferentially expressed in OAC or Barrett’s and performed GO term analysis (Supplementary Tables S8 and S9). Similar GO terms were identified with “MAPK signalling” and “Hallmark EMT/ECM organisation” (terms associated with cell migration) resembling those identified through association with eRNA regions (Fig. 4A). Similarly, Barrett’s-specific genes returned GO terms such as “epithelial cell differentiation”, as identified from eRNA regions further emphasising the similarity in biological processes identified by eRNA regions and total RNAs. To determine whether these similar GO categories reflected similar genes being identified, we overlapped the differentially expressed genes (DEGs; Supplementary Tables S8 and S9) with genes associated with differentially expressed eRNAs (DEEs; Supplementary Tables S2 and S3), that are enriched in either Barrett’s or OAC samples. We found a significant overlap between these sets of genes although the majority of the genes were uniquely identified by investigating either by total RNA-seq or by eRNA profiling (Fig. 4B). Therefore, despite pinpointing similar biological processes, eRNA profiling reveals different candidate genes involved these processes.

**Figure 4.**
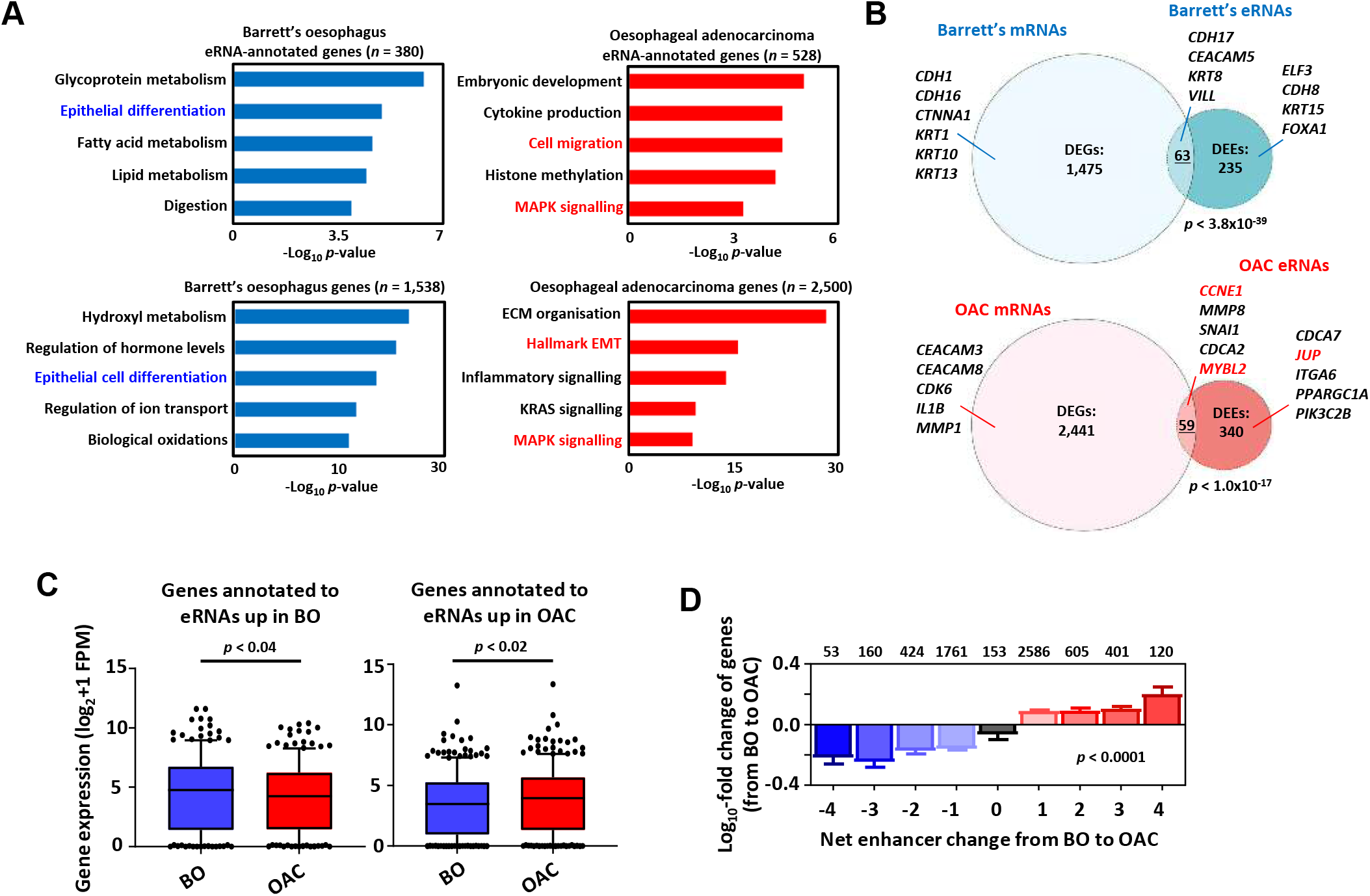
Association of eRNA regions with transcriptional regulators and potential target genes. (A) GO-term analysis of differentially expressed eRNA region-associated genes (top-left/right) and differentially expressed genes (bottom-left/right) in (left) Barrett’s (> Log_2_FC 0.9, < padj = 0.05) and (right) OAC (> Log_2_FC 1.1, < padj = 0.05). eRNAs were annotated to genes by the nearest gene model using HOMER (Heinz et al., 2010). (B) Venn diagram displaying overlap between differentially expressed genes and unique protein-coding genes annotated to differentially expressed eRNAs in (top) BO and (bottom) OAC (*p-*value is shown; Fisher’s exact test). (C) Box plots comparing the expression of genes annotated to eRNAs differentially expressed in BO (left) or OAC (right) in BO and OAC patient tissue total RNA-seq samples from the OCCAMS dataset. (p-value is shown; Welch’s t-test). (D) Genome-wide analysis of the effect of changing eRNA expression on gene expression within 200 kb chromosomal bins. Numbers above bars represent total genes associated with respective net-enhancer change (p-value is shown; Kruskal-Wallis test).

To further investigate whether the activities of enhancer regions are linked to nearby gene expression, we selected eRNA regions preferentially expressed in either OAC or Barrett’s and found that the nearest genes exhibited higher expression in the correct corresponding tissue type in two independent datasets (Fig. 4C; Supplementary Fig. S6A). This observation was further supported by comparing the expression of the genes closest to eRNA regions found in patient RNA-seq data to the expression of a random set of genes. This revealed significantly enhanced expression levels of eRNA-annotated genes in patients (Supplementary Fig. S6B). While the nearest gene model often correctly associates enhancers with the closest gene, this is not always the case, so we considered all genes within a 200 kb bin around the eRNA region rather than just the nearest gene. We then determined the net eRNA expression change when comparing Barrett’s to OAC samples and created 9 bins reflecting the magnitude of differential expression. We then calculated the associated gene expression changes within these genomic bins when comparing Barrett’s to OAC. There was a clear correlation between the directionality of eRNA expression with mRNA expression which changed in an analogous manner, with high eRNA levels in OAC associated with higher gene expression in OAC and vice versa in Barrett’s (Fig. 4D).

In summary therefore, eRNA expression profiling can reveal specific upstream regulatory transcription factors and the eRNA generating regions can be used to uncover a set of biological processes and constituent genes that are relevant to specific disease states.

### Target genes of eRNA-defined enhancers are co-expressed in OAC

We identified potential target genes of eRNA-defined enhancers by implementing the nearest gene model (Fig. 4B). However, the nearest gene is not always the enhancer target (Sanyal et al, 2012). We therefore examined the correlation between eRNA expression and the expression of their designated target genes for three candidate enhancers, localised in the vicinity of the *JUP* (Fig. 5A), *MYBL2* and *CCNE1* (Supplementary Fig. S7A and B) loci. Each of these putative enhancer regions contains more RNA signal in OAC compared to Barrett’s as well as evidence for chromatin accessibility in OAC patient material. We focussed on *JUP* as this had not been implicated in OAC previously and is significantly co-amplified with *ERBB2*, a key oncogenic driver of OAC (The Cancer Genome Atlas, 2017; Frankell et al., 2019; Supplementary Fig. S7C). Indeed, both *JUP* transcript and eRNA are upregulated compared to Barrett’s in OAC patients with high ERBB2 expression (Fig. 5B). We performed a similar analysis for *CCNE1* and *MYBL2* but instead examined their expression across all OAC samples. For both loci, the eRNA and gene transcript are both upregulated in OAC relative to Barrett’s (Supplementary Fig. S7D and E). Next, we examined the correlation of eRNA and transcript expression on a “sample by sample” basis. *JUP* eRNA expression showed strong correlation with *JUP* expression, irrespective of *ERBB2* level sample status (Fig. 5C). Lower correlations were observed with the expression of the two adjacent genes (Supplementary Fig. S7F) consistent with *JUP* being the relevant target. Similarly, *CCNE1* and *MYBL2* eRNA expression is more strongly correlated with the expression of their designated targets than either of their immediately flanking genes (Supplementary Fig. S7G and H). We extended this analysis across all eRNAs and asked whether we could find significantly correlated mRNA expression of genes located in their vicinity (+/-100kb)(Supplementary Table S10). Generally, highly correlated genes could be identified (see diagonal in Fig. 5D). Interestingly when we clustered the data according to the expression of the genes associated with each eRNA, then there was generally a good segregation of OAC-specific and BO-specific eRNAs further emphasising the relevance of the correlations we observed (Fig. 5D). Furthermore, when we split the RNAseq data into tissue types, there was a significantly higher correlation of BO-specific eRNA with nearby gene expression in BO datasets, compared to analysing the shared eRNAs (Fig. 5E, left). Similarly, the same trend was observed for OAC-specific eRNAs which were more highly correlated with nearby gene expression in OAC datasets (Fig. 5E, right).

**Figure 5.**
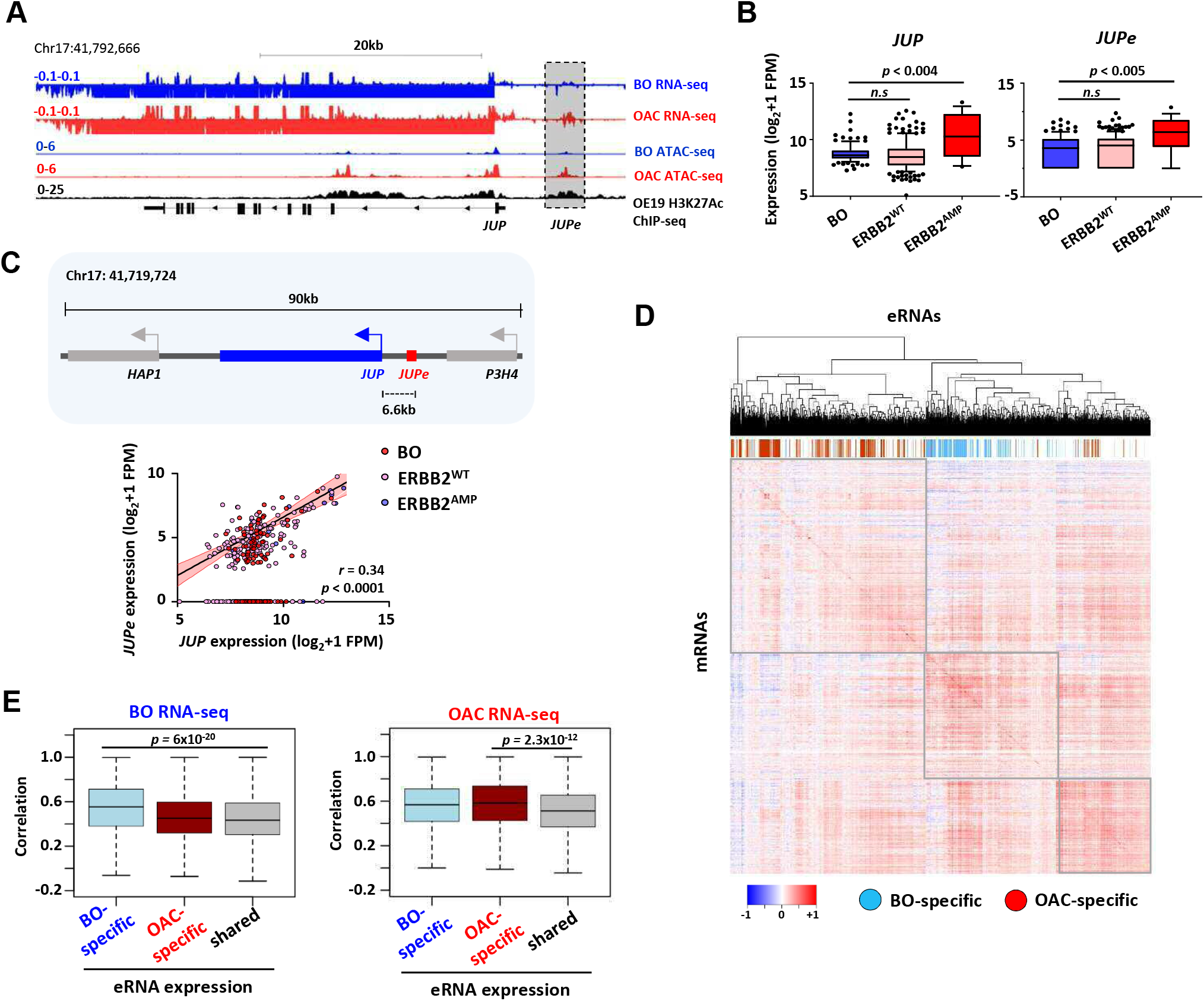
eRNA regions identify *JUP* as a candidate target gene. (A) Genome browser view of BO and OAC patient tissue ATAC- and total RNA-seq data, and H3K27ac ChIP-seq in OE19 cells, at the *JUP* locus with the *JUPe* eRNA highlighted. (B) Box plots comparing the expression of (left) *JUP* and (right) *JUPe* in BO (n = 108), ERBB2^WT^ (n = 193) and ERBB2^AMP^ (n = 17) OAC patient tissue total RNA-seq samples (p-value is shown; Welch’s t-test). (C) Schematic displaying relative locations of putative eRNA region target genes and nearest neighbours (top) and correlation of *JUPe* with *JUP* expression across BO (n = 108), ERBB2^WT^ (n = 193) and ERBB2^AMP^ (n = 17) OAC patient tissue total RNA-seq samples (Spearman’s r and p-value are shown; Spearman’s rank correlation test). (D and E) Global analysis of correlations of eRNA expression with the expression of the most correlated gene within a 200 kb window flanking the eRNA region. eRNAs are defined as tissue-specific according to Fig. S1A, and the rest of the eRNAs are designated as shared. (D) Heatmap showing the correlation coefficients between all 4600 eRNAs and associated mRNAs in the RNA-seq datasets. Samples are clustered according to eRNA-associated gene expression values. OAC-specific eRNAs (red), BO-specific eRNAs (blue) and shared eRNAs (white) are indicated across the top. (E) Boxplots showing the correlations with BO gene expression datasets (left) or OAC gene expression datasets (right). Significance values (t-test) are shown between the indicated groups.

Together these results demonstrate that we are able to link eRNA expression to their putative targets in the relevant disease-specific datasets.

### Validation of enhancer activity of eRNA regions

To validate that the regions generating eRNAs have enhancer activity we again focussed on the *JUP, MYBL2* and *CCNE1* loci. First, we showed that all three putative enhancer regions have significantly higher levels of KAS-seq signal in OE19 cells relative to a control enhancer from the *APOL4* gene that is not expressed in OE19 cells (Fig. 6A). This is reflective of ongoing transcription. Furthermore, all three eRNAs and their associated target genes exhibit higher expression in OE19 cells compared to the Barrett’s CP-A cell line (Fig. 6B). To directly establish enhancer activity, we cloned the regions encompassing the eRNAs into two different enhancer reporter systems with either luciferase (Fig. 6C) or RNA (Fig. 6D) readouts. For all three regions, both assays demonstrated significant enhancer activity in OE19 cells (Fig. 6C and D). Finally, we used an inducible dCas9-KRAB synthetic repressor protein to silence the activities of each enhancer in their natural chromatin context in OE19 cells (Supplementary Fig. S8A). In all cases, introduction of the relevant gRNA to target the dCas9-KRAB repressor to the putative enhancer, resulted in reduced eRNA transcription and reduced expression of the associated target gene (Fig. 6E). Importantly, no significant changes in expression were observed for any of the genes immediately flanking the target genes, demonstrating the fidelity of our enhancer-gene linkages (Supplementary Fig. S8B).

**Figure 6.**
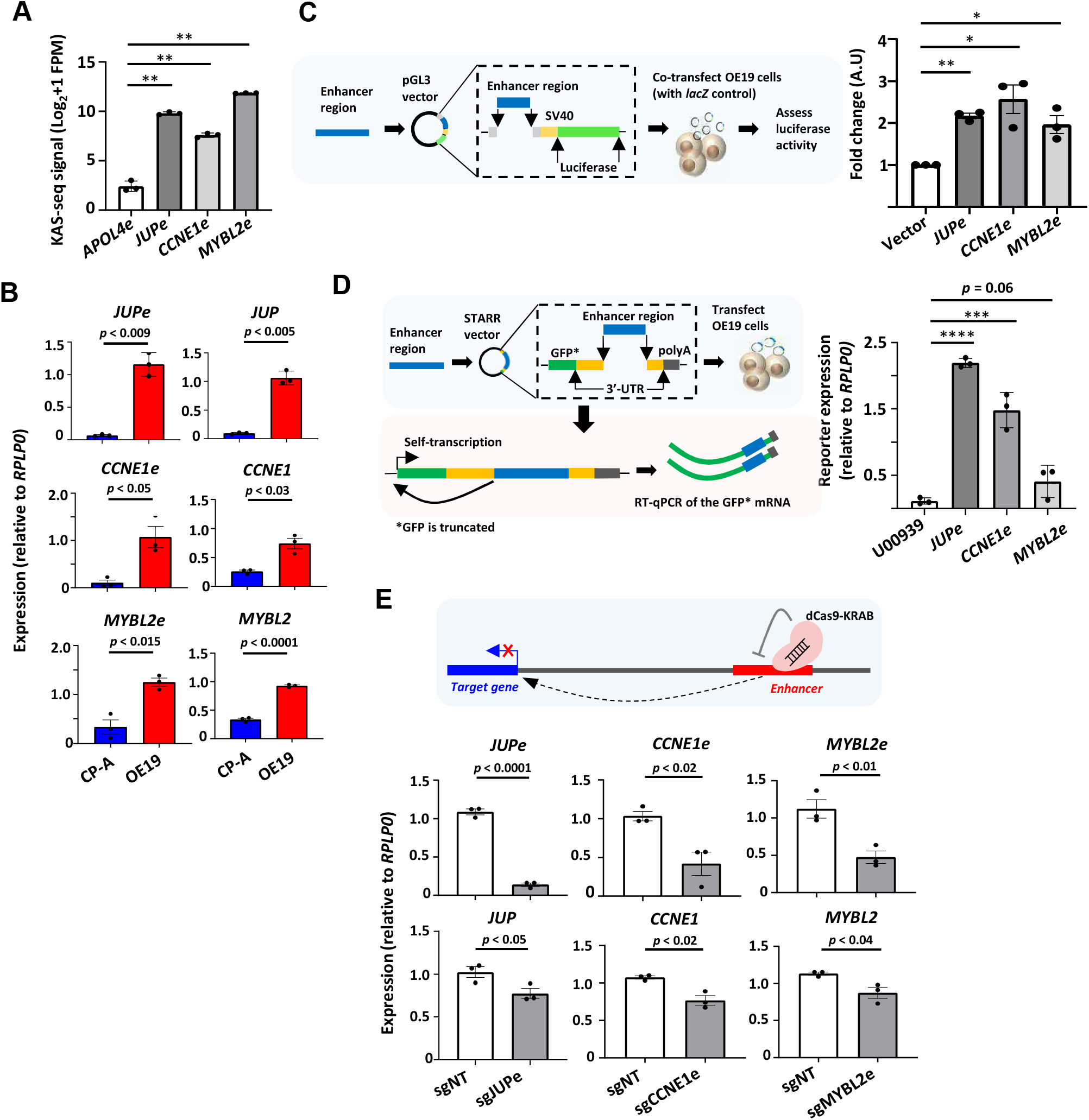
*In vitro* interrogation of eRNA regions confirms production and association with cancer-associated processes. (A) Bar graphs displaying KAS-seq signal at the *APOL4e, JUPe, CCNE1e* and *MYBL2e* regions in OE19 cells (** = *p* < 0.01; Welch’s t-test). (B) Bar graphs displaying difference in expression of *JUP, CCNE1, MYBL2* and *JUPe, CCNE1e*, and *MYBL2e* between CP-A and OE19 cells using RT-qPCR (*p*-value is shown; Welch’s t-test). (C) (Left) Schematic of luciferase assay. Bar graph displaying the difference in luciferase reporter activity between *JUPe, CCNE1e* and *MYBL2e*, compared to vector only negative control (** = *p* < 0.01; * = *p* < 0.05; one-way ANOVA with Bonferroni’s correction). (D) (Left) Schematic of STARR-RT-qPCR assay. (Right) Bar graph displaying the difference in STARR reporter activity between *JUPe, CCNE1e* and *MYBL2e*, compared to *U00930* tRNA negative control (**** = *p* < 0.0001; *** = *p* < 0.001; one-way ANOVA with Bonferroni’s correction). (E) Bar graphs displaying the expression of *JUPe, CCNE1e, MYBL2e* eRNAs (top) and *JUP, CCNE1* and *MYBL2* mRNAs (bottom) in OE19-dCas9-KRAB cells using real time RT-qPCR, upon treatment with the indicated targeting or non-targeting (NT) sgRNA (*p*-value is shown; Welch’s t-test). A schematic of dCas9-KRAB targeting of eRNA regions is shown.

Together these results build on our correlative observations linking eRNA containing regions with enhancer like properties and provide definitive proof of enhancer activity and regulatory linkage to neighbouring genes.

### Biological and clinical relevance of eRNAs and their target genes

We have shown that the discovery of eRNAs in OAC patients reveals genes and processes which are operative in OAC and allows us to distinguish OAC from Barrett’s patients. To provide further biological insights, we asked whether any of the three eRNA target genes, *JUP, MYBL2* and *CCNE1* were uncovered in a cell line viability screen in the DepMap project (Tsherniak et al., 2017; Behan et al., 2019). We found that four of the top six cell lines showing a dependency on *JUP* expression are gastroesophageal in origin and these all contain *ERBB2* amplifications (Fig. 7A, left). *JUP* is also the highest scoring gene for fitness dependency across OAC cell lines (Fig. 7A, right). However, *MYBL2* and *CCNE1* did not score highly in this screen. We therefore further probed the function of the eRNA-defined enhancers in the OE19 OAC cell line by using the dCas9-KRAB silencing system directed at these regions. In all cases, enhancer silencing led to significant reductions in cell viability and growth (Fig. 7B; Supplementary Fig. S9A).

**Figure 7.**
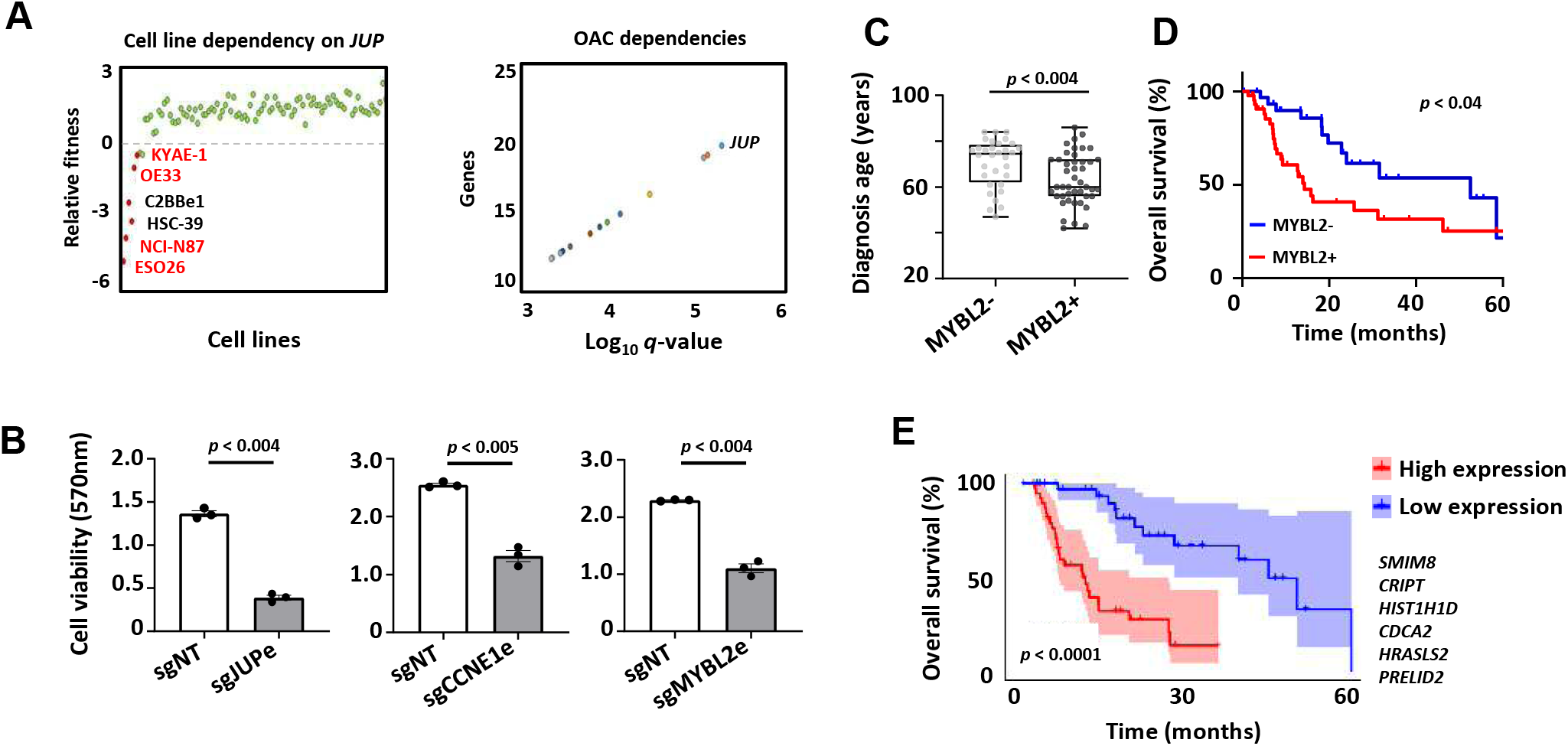
Biological and clinical relevance of eRNAs and their target genes. (A) Scatter plots displaying data from the Sanger DepMap Project Score (Tsherniak et al., 2017; Behan et al., 2019) highlighting (left) cell line dependency on *JUP* (gastroesophageal cell lines are marked in red) and (right) top genetic dependencies in OAC. (B) Bar graph displaying the difference in cell viability in OE19-dCas9-KRAB cells upon sgRNA treatment, assessed by crystal violet assay (*p*-value is shown; Welch’s t-test). (C) Box plots comparing diagnosis age for OAC patients with low and high *MYBL2* expression in the TCGA PanCancer Atlas dataset (*p*-value is shown; Welch’s t-test). (D) Kaplan-Meier plot comparing overall survival between OAC patients with low and high *MYBL2* expression in the TCGA PanCancer Atlas dataset (Log rank *p*-value is shown). (E) Kaplan-Meier plot comparing overall survival between OAC patients with low and high signature eRNA target expression in the TCGA PanCancer Atlas dataset (Log rank *p*-value is shown; signature genes are shown).

To provide further clinical relevance, we used an RNA-seq dataset from the TCGA consortium (The Cancer Genome Atlas, 2017) that differed from our discovery cohort to ask whether any of the eRNA target genes informed on any particular clinical features. We found that the age of diagnosis was lower in patients expressing high levels of *MYBL2* (Fig. 7C) and *JUP* (Supplementary Fig. S9B) suggesting earlier disease onset. Furthermore, in the case of *MYBL2*, high level expression was indicative of lower median survival times (Fig. 7D), although *JUP* and *CCNE1* were not informative in that regard (Supplementary Fig. S9C). Altogether, 32% of genes annotated to OAC-specific eRNAs displayed a significant prognostic value for patient survival (Supplementary Fig. S9D). Finally, we took an unbiased approach and asked whether we could identify a signature within the eRNA-associated genes which had clinical significance. This revealed a 6-gene signature that was highly predictive of OAC patient survival (Fig. 7E).

Collectively, these data demonstrate the functional importance of the eRNA-defined enhancers and their target genes for OAC cell growth and their potential utility for assessing patient prognosis. In the case of *JUP*, the broad OAC cell dependency suggests that this represents a target of potential therapeutic value, especially in *ERBB2*-positive patients.

## Discussion

Cancer is driven by a combination of genetic and epigenetic changes (reviewed in Zhao et al., 2021). Both of these processes ultimately lead to alterations in the activity of gene regulatory elements, including transcriptional enhancers, that results in a change in cellular phenotype that defines the tumourigenic state. While profiling of histone marks and chromatin accessibility is useful in defining potential gene regulatory elements, this approach is limited for defining active enhancers. Here we used eRNA profiling to identify regions harbouring potentially active enhancers in oesophageal adenocarcinoma patient samples. We integrated these with a range of epigenetic datasets and experimentally validated several regions as bona fide enhancers. Importantly, our enhancer repertoire identified new pathways that are activated in OAC which were not apparent from either genome sequencing or mRNA profiling alone.

A previous pan-cancer analysis of RNA-seq data sets generated by the TCGA consortium to identify eRNAs defined a compendium of potential enhancers across human cancers and demonstrated how they could have clinical significance (Chen et al., 2018). However, while the authors examined oesophageal cancers, they mixed two distinct disease sub-types, squamous and adenocarcinoma, which limited any discoveries specific to OAC. Here we specifically interrogated OAC RNA-seq data (generated by the OCCAMs consortium) and to identify enhancers that are potentially relevant to OAC, we compared their associated eRNA levels to the pre-cancerous Barrett’s oesophagus state. Using this approach, we were able to identify ∼1000 high confidence OAC-specific enhancers. These enhancer regions exhibited high accessibility in both patient samples and cell line models and using a variety of chromatin marks profiled in an OAC cell line model we provided further verification of the enhancer-like properties. The OAC-specific enhancers are associated with transcription factors which have been shown to be important for driving OAC-specific transcriptional events (eg KLF5, Rogerson et al., 2020; AP1, Britton et al., 2017). Reciprocally we also identify ∼700 Barrett’s specific enhancers which are associated with a different transcription factor repertoire, including the potential involvement of members of the TP53/TP63/TP73 family. While we have identified a large number of intergenic enhancers, the approach we have taken will miss intragenic enhancers, and other approaches using function-based assays (eg STARR-seq; Arnold et al., 2013) or computational imputation will be needed to identify these.

Our newly derived eRNA defined enhancer data sets also provide novel insights into pathways that are operational in OAC. This is apparent from the limited overlap in differentially expressed genes at the mRNA level versus the differential expression of genes associated with nearby enhancers defined by eRNA levels. Interestingly while the specific gene overlaps are limited, the broad processes defined by GO terms such as MAPK signalling and cell migration/EMT remain the same. Part of this discrepancy might be explained by the OAC-specific enhancers maintaining gene expression in the BO-OAC transition rather than *de novo* gene activation in OAC. However, expression change cut offs we use may also contribute to this, as can the heterogeneity of the OAC samples. An enhancer associated with *JUP* was specifically revealed by eRNA profiling, alongside hundreds of other enhancers linked to genes involved in oncogenic processes such as cell migration, PI3K signalling and metabolism. We validated this *JUP* enhancer, and enhancers linked to *MYBL2* and *CCNE1*, and their association with their proposed targets by CRISPRi. Furthermore, correlations between eRNA and mRNA expression across cancer samples suggest a causal link. Importantly both CCNE1 and MYBL2 play important roles in promoting cell proliferation, an important cancer cell trait. JUP (otherwise known as junction plakoglobin) had not previously been implicated in OAC but this was identified in a screen for gene dependencies in OAC cell lines (DepMap project: Tsherniak et al., 2017; Behan et al., 2019) and we validated its importance for OAC cell growth. In this context, the co-amplification with ERBB2 is intriguing as both genes are on the same chromosome and are ∼2 Mb apart and the intervening region is not usually co-amplified. This may reflect a functional interdependency for these two oncogenic events. JUP has previously been implicated in multiple cancers although is generally found to be a tumour suppressor protein, rather than the oncogenic properties it has in the context of OAC (reviewed in Aktary et al., 2017). As JUP encodes a protein involved in cell-cell contacts, this might suggest a role for this process in OAC cancer cell survival and a potential route to therapy. Alternatively, JUP may be acting via the numerous other cellular processes in which it has been implicated, and further work is needed to understand the precise role it has in OAC cells.

In addition to pointing to potential actionable pathways, we also demonstrate that eRNA profiling is clinically relevant and is sufficient to differentiate between BO and OAC. A six gene signature derived from our OAC-specific enhancer associated genes is able to predict prognostic outcomes. Indeed, a large proportion of the eRNA associated genes show prognostic significance when analysed on an individual basis. Furthermore, by focussing in on a few examples, we found that one of the novel OAC-associated genes, *JUP*, was upregulated in *ERBB2* overexpressing OAC samples which is reflected by their frequent co-amplification. Coupled with the observation that JUP is required for the survival of a range of OAC cell lines harbouring *ERBB2* amplifications, this further emphasises the potential utility of JUP as a therapeutic target in this subset of OAC patients. This would provide an alternative approach to the use of ERBB2 inhibitors which are routinely administered but have limited therapeutic benefit (Bang et al., 2010). Further clinical insights are provided by other eRNA defined enhancer regions, such as the enhancer associated with *MYBL2* where high *MYBL2* expression indicates a worse prognosis for patients and earlier disease onset.

In summary, we identify a cohort of OAC-specific enhancers, expanding our knowledge of the regulatory networks that are operational in OAC. This has led to novel insights into the pathways that are operational in this disease. The approach we have taken to identify cancer-specific enhancers should be broadly applicable to other tumour types or subtypes, where data are available for both the cancer and the originating normal or pre-cancerous tissue.

## Materials and methods

### Cell culture and treatments

OE19 cells were cultured in RPMI 1640 (ThermoFisher Scientific, 52400) supplemented with 10% foetal bovine serum (ThermoFisher Scientific, 10270). OE19-dCas9-KRAB stable cells were previously generated (Rogerson et. al, 2020) and cultured as above with the addition of 500 ng/mL puromycin (Sigma P7255). The expression of dCas9-KRAB was induced using 250 ng/mL doxycycline (Sigma-Aldrich, D3447. Cell lines were cultured at 37°C, 5% CO_2_ in a humidified incubator.

### Dominant negative FOS over-expression

pINDUCER20-GFP-AFOS (ADS5006, Britton et al., 2017) was packaged into lentivirus and OE19 cells were transduced with lentivirus as previously described (Tiscornia et al., 2006). Briefly, 3×10^6^ HEK293T cells were transfected with 2.25 µg psPAX2 (Addgene, 12260), 1.5 µg pMD2.G (Addgene, 12259) and 3 µg pINDUCER20-GFP-AFOS using PolyFect (Qiagen, 301107). Media was collected at 48 and 72 hours post-transfection and viral particles were precipitated using PEG-it™ Solution (System Biosciences, LV810A-1). To transduce, cells were treated with virus (MOI 0.5-1.0) and 5 µg/mL Polybrene (EMD Millipore, TR-1003). Polyclonal cells were selected for 2 weeks in 250 µg/mL G418 (ThermoFisher Scientific, 10131027). Dominant negative FOS (dnFOS; Olive et al., 1997) was induced with 1 µg/mL doxycycline.

### sgRNA transfection

2×10^5^ cells were transfected with 10 pmol sgRNA pool using Lipofectamine™ RNAiMAX transfection reagent (ThermoFisher Scientific, 13778150) according to the manufacturer’s instructions. Cells were seeded into 6-well plates. Modified full length sgRNAs were designed using E-CRISP (Heigwer et al., 2014) and off-target activity assessed using CCTop (Stemmer et al., 2015). sgRNAs were ordered from Synthego. sgRNA sequences are listed in Supplementary Table S11.

### Cell growth and cell viability assays

Cell growth and viability was assessed by crystal violet assay. Assays were performed by fixing cells in 4% paraformaldehyde for 10 minutes. Cells were stained with 0.1% crystal violet (Sigma-Aldrich, HT90132) for 30 minutes. Crystal violet dye was extracted using 10% acetic acid and absorbance readings taken at 570 nm on a SPECTROstar Nano Microplate Reader (BMG LABTECH). Cell growth measurements were taken at 0, 24, 48 and 72 hours and cell viability measurements taken at 72 hours.

### RT-qPCR and eRNA qPCR

Total RNA was extracted from cells using an RNeasy Plus RNA extraction kit (Qiagen, 74136) according to the manufacturer’s protocol. RT-qPCR reactions were run using the QuantiTect SYBR Green RT-qPCR kit (Qiagen, 204243) on a Qiagen Rotor-Gene Q. For eRNA-qPCR, RNA was extracted using an RNeasy Plus RNA extraction kit (Qiagen, 74136) with the on-column DNAse digest, according to manufacturer’s instructions. 500ng of RNA was reverse-transcribed using SuperScript™ VILO™ Master Mix (ThermoFisher Scientific, 11755250) according to manufacturer’s instructions. eRNA levels were assessed by qPCR using a Rotor-Gene SYBR Green PCR Kit (Qiagen, 204074) on a Qiagen Rotor-Gene Q. Relative copy number of transcripts was determined by standard curve and normalised to the expression of *RPLP0* control gene. Primers used are listed in Supplementary Table S11.

### Luciferase and STARR-qPCR reporter assays

Regions containing JUPe, MYBL2e or CCNE1e were amplified from OE19 genomic DNA using primers containing 20 bp overlap regions with the multiple cloning site of the pGL3 Promoter vector (Promega, E1761) for luciferase assays, or between the InFusion arms of the hSTARR ORI vector (Addgene, 99296) (Supplementary Table S11. Final vectors were assembled using HiFi assembly (NEB, E5520S) according to the manufacturer’s instructions to create plasmids containing JUPe, MYBL2e or CCNE1e enhancer regions in either hSTARR ORI (pAS5008-pAS5010) or pGL3-vectors (pAS5011-pAS5013). Enhancer vectors were transfected using the Amaxa™ Nucleofector™ II (Lonza) with Cell Line NucleofectorTM Kit V (Lonza, VCA-1003) and program T-020, according to manufacturer’s instructions. For luciferase assays, 250ng of enhancer vector was co-transfected alongside 50ng of pCH110 (Amersham). For STARR-qPCR, 800ng of vector was transfected. Enhancer activity was assessed using the Dual-Light™ Luciferase & β-Galactosidase Reporter System (ThermoFisher Scientific, T1003) according to the manufacturer’s instructions, or by RT-qPCR.

### Western blots

Cells were lysed in RIPA buffer (1% IGEPAL CA-630, 150 mM NaCl, 0.1% SDS, 50 mM Tris pH 8.0, 1 mM EDTA, 0.5% sodium deoxycholate) and protease inhibitor cocktail supplement (Roche, 11836170001). Protein concentration was determined by bicinchoninic acid assay (Pierce, 23227). 5x SDS loading buffer (235 mM SDS, 10% -mercaptoethanol, 0.005% bromophenol blue, 210 mM Tris-HCl pH 6.8, 50% glycerol) was added to lysates to a final 1x concentration and incubated for 10 minutes at 90°C. Proteins were then resolved by SDS-PAGE and transferred onto a nitrocellulose membrane. Membranes were blocked using Odyssey® Blocking Buffer (LI-COR Biosciences, P/N 927-40000). Antibodies used: anti-Cas9 (Diagenode, C15200229, 1:10,000) and anti-ERK (Cell Signaling Technologies, 4695S, 1:1,000). Secondary antibodies used: anti-rabbit (LI-COR Biosciences, 926-32213, 1:10,000) and anti-mouse (LI-COR Biosciences, 926-32210, 1:10,000). Membranes were visualized using a LI-COR Odyssey® CLx Infrared Imager

### eRNA and mRNA analysis

Patient tissue ATAC-seq data processing was performed as described previously (Britton et al., 2017). Reads were mapped to GRCh38 (hg38) using Bowtie2 v2.3.0 (Langmead and Salzberg, 2012) with the following options: -X 2000 -dovetail. Mapped reads (>q30) were retained using SAMtools (Li et al., 2009). Reads mapping to blacklisted regions were removed using BEDtools (Quinlan and Hall, 2010). Peaks were called using MACS2 v2.1.1 (Zhang et al., 2008) with the following parameters: -q 0.01, -nomodel-shift -75 -extsize 150 -B -SPMR. A custom union peakset was formed from all BO and OAC patient samples, using HOMER v4.9 mergePeaks.pl -d 250 (Heinz et al., 2010) as described previously (Rogerson et al., 2019) and filtered to retain only intergenic regions :2 kb upstream from a TSS or ≥500 bp downstream from a TTS.

RNA-seq reads were mapped to the human genome GRCh38 (hg38) using STAR v2.3.0 (Dobin et al., 2013). Expression threshold for eRNAs was determined using an adapted method from Zhang et al., 2019 (Zhang et al., 2019). Briefly, total RNA-seq reads were integrated into genomic regions from the intergenic patient ATAC-seq peakset. Putative eRNA and mRNA read counts were determined using featureCounts (Liao et al., 2014) and FPM values determined using DESeq2 (Love et al., 2014). Putative eRNA regions with average counts and FPM values of ≥3 and 1.5, respectively, were taken forward for further analysis. Differentially expressed eRNAs and mRNAs were determined using DESeq2 (Love et al., 2014). For eRNAs, a log_2_-fold change of ±0.5 and P-value_adj_ < 0.05 defined differential expression. For BO and OAC mRNAs, a log_2_-fold change of ±0.9 and ±1.5, respectively, and P-value_adj_ < 0.05 defined differential expression. ERBB2-positive OAC samples (*ERBB2*^AMP^) were determined based on these samples having expression of *ERBB2* greater than the median *ERBB2* expression +2 SD. Morpheus (https://software.broadinstitute.org/morpheus/) was used to generate heatmaps and perform hierarchical clustering.

HOMER v4.9 was used for de novo transcription factor motif enrichment analysis. To analyse footprinting signatures at putative eRNA regions, TOBIAS v0.5.1 was used (Bentsen et al., 2020). eRNAs were annotated to genes by the nearest gene model and assessed for CpG content using HOMER v4.9. Super enhancers were identified using HOMER v4.9 findPeaks.pl -style super. Net enhancer activity was calculated as in Bi et al., 2020 (Bi et al., 2020). Briefly, neighbouring genes of eRNA regions in both BO and OAC were identified and stratified into nine groups based on the net eRNA change within 200 kb of the TSS of each gene: + (or -1) stands for 1 net gained (or lost) eRNA from BO to OAC. Bidirectionality score was calculated using HOMER v4.9 analyzeRepeats.pl with the -strand option applied for each strand and score defined as log_10_((+strand expression score + 1)/(-strand expression score + 1)) + 1.

### DepMap data

Batch-corrected genome-wide CRISPR-Cas9 knockout screen data (DepMap Public 21Q4 CRISPR gene dependency.csv) was obtained from DepMap (https://depmap.org/portal/).

### ChIP-seq data analysis

ChIP-seq analysis was carried out as described previously (Wiseman et al., 2015). OE19 H3K27ac and GAC H3K4me1/3 ChIP-seq reads were mapped to the human genome GRCh38 (hg38) using Bowtie2 v2.3.0 (Langmead and Salzberg, 2012). Biological replicates were checked for concordance (r > 0.80). Peaks were called using MACS2 v2.1.1, using input DNA as control (Zhang et al., 2008). Mapped reads (>q30) were retained using SAMtools (Li et al., 2009). Reads mapping to blacklisted regions were removed using BEDtools (Quinlan and Hall, 2010).

### dnFOS RNA-seq processing and data analysis

dnFOS was induced in OE19-dnFOS cells for 48 hours before RNA was isolated and sequenced. Three biological replicates were sequenced per condition. Total RNA was extracted from cells using an RNeasy Plus RNA extraction kit (Qiagen, 74136). The on-column DNase digest (Qiagen, 79254) was performed according to the manufacturer’s protocol. RNA-seq libraries were generated using a TruSeq stranded mRNA library kit (Illumina, RS-122-2001) and sequenced on an Illumina HiSeq 4000 System (University of Manchester Genomic Technologies Core Facility).

Reads were mapped aligned to the human genome GRCh38 (hg38) using STAR v2.3.0 (Dobin et al., 2013). Gene expression counts were obtained using featureCounts (Liao et al., 2014) and differentially expressed genes were identified by DESeq2 using P-value_adj_ < 0.05 (Love et al., 2014). OE19 siKLF5 RNA-seq (Rogerson et al., 2020) data analysis was also performed as above.

### CUT&TAG processing and data analysis

CUT&TAG library generation was performed as described previously (Kaya-Okur et al., 2020) with an altered nuclear extraction step. For the nuclear extraction, OE19 cells were initially lysed in Nuclei EZ lysis buffer (Sigma-Aldrich, NUC-101) at 4°C for 10 mins followed by centrifugation at 500 g for 5 mins. The subsequent clean-up was performed in a buffer composed of 10 mM Tris-HCl pH 8.0, 10 mM NaCl and 0.2% NP40 followed by centrifugation at 1300 g for 5 mins. Nuclei were then lightly cross-linked in 0.1% formaldehyde for 2 mins followed by quenching with 75 mM glycine followed by centrifugation at 500 g for 5 mins. Cross-linked nuclei were resuspended in 20 mM HEPES pH 7.5, 150 mM NaCl and 0.5 M spermidine at a concentration of 4-8×10^3^/µL (2-4×10^4^ total). Subsequent stages were as previously described (Kaya-Okur et al., 2020). For 2-4 ×10^4^ nuclei, 0.5 µg of primary and secondary antibody were used with 1 µL of pA-Tn5 (Epicypher, 15-1017). Antibodies used: anti-BRD4 (abcam, ab128874), anti-CTCF (Merck-Millipore, 07-729), anti-H3K27ac (abcam, ab4729), anti-H3K27me3 (Merck-Millipore, 07-449), anti-H3K4me1 (abcam, ab8895), anti-H3K4me2 (Diagenode, pAb-035-010), anti-H3K4me3 (abcam, ab8580), anti-H3K36me3 (Diagenode, pAb-058-010), anti-H4K20me1 (Diagenode, mAb-147-010), anti-PolII (abcam, ab817), anti-PolII-S2 (abcam, ab5095), anti-PolII-S5 (abcam, ab5131) and anti-Med1 (AntibodyOnline, A98044/10UG). CUT&TAG libraries were pooled and sequenced on an Illumina HiSeq 4000 System (University of Manchester Genomic Technologies Core Facility). CUT&TAG data processing was performed as for ChIP-seq but with the MACS2 v2.1.1 (Zhang et al., 2008) but the --broad peak calling option was used for the H4K20me1, H3K27me3 and H3K36me3 marks. Fraction reads in peak (FRiP) scores for each mark were calculated using featureCounts and a stringent threshold of ≥2% was set to ensure quality of data for downstream analyses (Landt et al., 2012; FRiP scores are listed in Supplementary Table S12). ChromHMM (Ernst and Kellis., 2012) was used to train an eight-state Hidden Markov Model using the CUT&TAG data for all marks assayed. The number of states was determined by running the model with increasing numbers of states until state separation was observed. Emission states were annotated in accordance with Roadmap Epigenomics Consortium Data (Roadmap Epigenomics Consortium et al., 2015).

### KAS-seq processing and data analysis

KAS-seq library generation was performed as described previously (Wu et al., 2020) except with nuclear extraction and labelling reactions. Nuclei were extracted and washed as described for CUT&TAG. Nuclei were then resuspended in nuclease-free H_2_O at a concentration of 1×10^4^/µL. (2×10^5^ total) Labelling reactions were carried out in DNA LoBind® tubes (Eppendorf, 0030108051) using 5 mM N_3_-kethoxal (a gift from Chuan He) in PBS to a final volume of 50 µL for 15 mins at 37°C with 1000 RPM mixing in a thermomixer. Labelled gDNA was isolated using the PureLink™ Genomic DNA Mini kit (ThermoFisher Scientific, K182001) and eluted twice with 21.5 µL 25 mM K_3_BO_3_ pH 7.0. Subsequent library preparation stages were as previously described (Wu et al., 2020). KAS-seq libraries were pooled and sequenced on an Illumina HiSeq 4000 System (University of Manchester Genomic Technologies Core Facility). Three biological replicates were sequenced and checked for concordance (r > 0.80). KAS-seq data processing was performed as described previously (Wu et al., 2020), but with the MACS2 v2.1.1 --broad peak calling option.

### The Cancer Genome Atlas data

Diagnosis age and overall survival between OAC patients with high or low *JUP, CCNE1* and *MYBL2* RNA expression (defined as ±1 SD from the median expression) in the TCGA PanCancer Atlas dataset (Liu et al., 2018) was downloaded from cBioPortal (https://www.cbioportal.org/study/summary?id=esca_tcga_pan_can_atlas_2018). Oncoprint plot of mutational co-occurrence between *JUP* and *ERBB2* in OAC was generated using cBioPortal.

To establish the prognostic model, Univariate Cox regression was performed using the survival package in R v3.6.0 to select genes associated with patient prognosis. A random forest algorithm was applied using the randomForestSRC package in R v3.6.0 for feature reduction to obtain a survival signature. Risk score (risk score = ∑*x*_i_ x β_i_ where *x*_i_ is gene expression value; β_i_ is coefficient index) was calculated using a Multivariate Cox regression model. Patients were grouped by the median value of risk score and Kaplan-Meier analysis performed to compare the survival difference between high and low risk score group. Visualisation was achieved using the survminer package in R v3.6.0.

### Bioinformatics

Genome browser data was visualised using the UCSC Genome Browser (Kent et al., 2002). Heatmaps and tag density plots of epigenomic data were generated the using deepTools (Ramirez et al., 2016) computeMatrix, plotProfile, plotCorrelation and plotHeatmap functions. Metascape (Zhou et al., 2019) was used for gene ontology analysis of gene sets. The eulerr package in R v3.6.0 was used for generating Venn diagrams.

### Datasets

All data was obtained from ArrayExpress, unless stated otherwise. Human tissue RNA-seq data was obtained from: OCCAMS consortium (European Genome-Phenome Archive, EGAD00001007496). Human tissue ATAC-seq data was obtained from: E-MTAB-5169 (Britton et al., 2017), E-MTAB-6751 (Rogerson et al., 2019), E-MTAB-8447 (Rogerson et al., 2020). The Cancer Genome Atlas OAC ATAC-seq data were obtained from the GDC data portal (portal.gdc.cancer.gov; Corces et al., 2018). OE19 H3K27ac ChIP-seq was obtained from: E-MTAB-10334 (Ogden et al., 2021). GAC H3K4me1 and H3K4me3 ChIP-seq was obtained from: Gene Expression Omnibus, GSE75898 (Ooi et al., 2016). OE19 siKLF5 RNA-seq and KLF5 ChIP-seq were obtained from: E-MTAB-8446 and E-MTAB-8568, respectively (Rogerson et al., 2020).

### Data access

All data have been deposited at ArrayExpress; OE19 KAS-seq and CUT&TAG data (E-MTAB-11357 and E-MTAB-11356, respectively) and OE19 dnFOS RNA-seq (E-MTAB-10334).

## Acknowledgements

We thank Guanhua Yan for excellent technical assistance, and staff in the Bioinformatics, and Genomic Technologies core facilities. We also thank Nicoletta Bobola and Sankari Nagarajan, for critical appraisal of the manuscript. We thank all lab members, Hannah Reed and Connor Rogerson for experimental and analytical advice. We are grateful to Chuan He for providing N_3_-kethoxal. This work was funded by grants to ADS from the MRC (MR/V010263/1) and the Wellcome Trust (102171/B/13/Z).

## Author contributions

I.A., S.O. and S-H.Y. performed the experiments and data analysis in this study; Y.L. and W.Z. performed data analysis; A.D.S. led the study. All authors contributed to manuscript preparation and/or critically appraised manuscript drafts.

