## Supplementary Figures for "eRNA profiling uncovers the enhancer landscape of oesophageal adenocarcinoma and reveals new deregulated pathways"

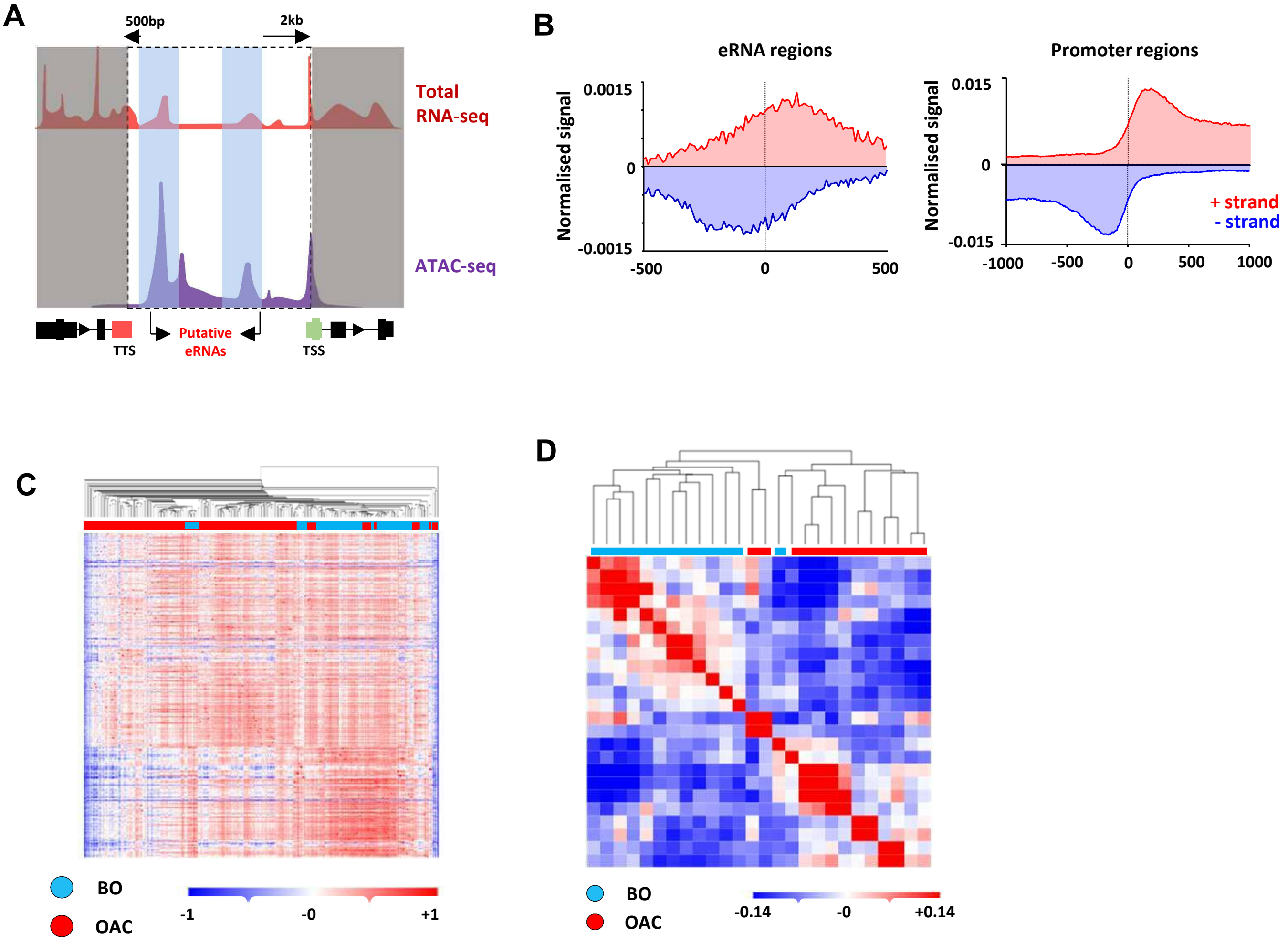


**Figure S1. Identification of enhancer transcription in BO and OAC patients.**
(A) Diagram of the sites of transcription considered for enhancer analysis. Total RNA-seq reads were integrated with intergenic regions of accessible chromatin at least 500 bp downstream/2 kb upstream of genes. (B) RNA-seq signal in BO *(n* = 108*)* and OAC *(n* = 210*)* patient tissue total RNA-seq samples plotted in a strand-specific manner across a 1 kb region centred on the eRNA containing regions (left) or a 2 kb region centred on the promoter regions (right). Data are normalised for total number of reads. (C) Pearson's correlation and hierarchical clustering of BO *(n* = 108*)* and OAC *(n* = 210*)* patient tissue total RNA-seq samples (OCCAMs) according to row z-score normalised gene expression levels. (D) Pearson's correlation and hierarchical clustering of BO *(n* = 13*)* and OAC *(n* = 13*)* patient tissue total RNA-seq samples (Maag et al., 2017 according to row z-score normalised expression levels in the 4600 eRNA regions.

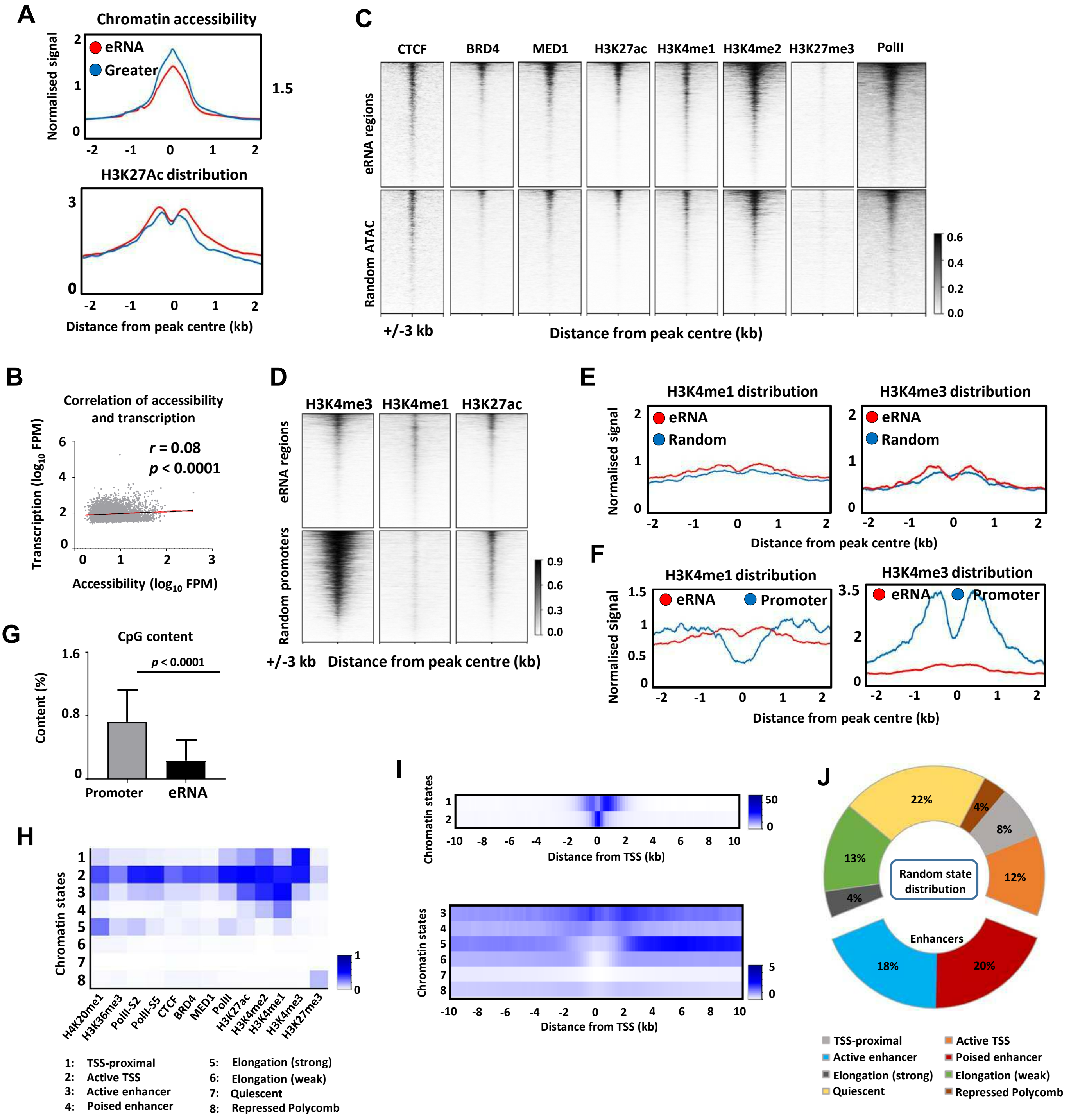


**Figure S2. Putative eRNAs are associated with enhancer-like genomic regions.**
(A) Metaplots of patient tissue chromatin accessibility and OE19 cell H3K27ac ChIP-seq signal at 4,600 eRNA regions, compared to 4,600 random regions of accessible chromatin of greater accessibility level. (B) Correlation of accessibility and transcription at 4,600 eRNA regions in BO ($n$ = 4) and OAC ($n$ = 14) patient tissue ATAC-seq samples and BO ($n$ = 108) and OAC ($n$ = 210) patient tissue total RNA-seq samples (Spearman's $r$ and $p$-value are shown; Spearman's rank correlation test). (C) Heatmaps showing CUT&TAG signal in OE19 cells at (top panels) 4,600 eRNA regions compared against (bottom panels) 4,600 randomly selected regions of accessible chromatin. Heatmaps for BRD4, CTCF, MED1, H3K27ac, H3K4me1, H3K4me2 and H3K27me3 are shown. (D Heatmaps showing (left) CUT&TAG signal in OE19 cells at (top panels) 4,600 eRNA regions compared against (bottom panels) 4,600 randomly selected accessible promoters. Heatmaps for H3K36me3, H3K4me3, H3K4me1, H3K27ac and H3K27me3 are shown and heatmaps showing (right) CUT&TAG signal in OE19 cells at (top panels) 4,600 eRNA regions compared against (bottom panels) 4,600 randomly selected regions of accessible chromatin. Heatmaps for PolII, PolII-S2, PolII-S5, and H3K4me3 are shown. (All heatmap regions are shown as peak centre ± 3 kb). (E) Metaplots of (left) H3K4me1 and (right) H3K4me3 GAC patient tissue ChIP-seq signal at 4,600 eRNA regions compared to 4,600 randomly selected region of accessible chromatin. (regions shown as peak centre ± 2 kb). (F) Metaplots of (top-right) H3K4me1 and (bottom-right) H3K4me3 GAC patient tissue ChIP-seq signal at all 4,600 eRNA regions compared to 4,600 randomly selected accessible promoters (all regions shown as peak centre ± 2 kb). (G) Bar graph displaying CpG content at eRNAs vs randomly selected accessible promoters ($p$-value is shown; Kolmogorov-Smirnov test). (H) Heatmap of chromatin states generated from CUT&TAG data using ChromHMM. (I) Heatmaps displaying distance from TSS for chromatin states (top) 1 and 2, and (bottom) 3-8. (J) Distribution of ChromHMM chromatin states for 4,600 random regions of accessible chromatin.

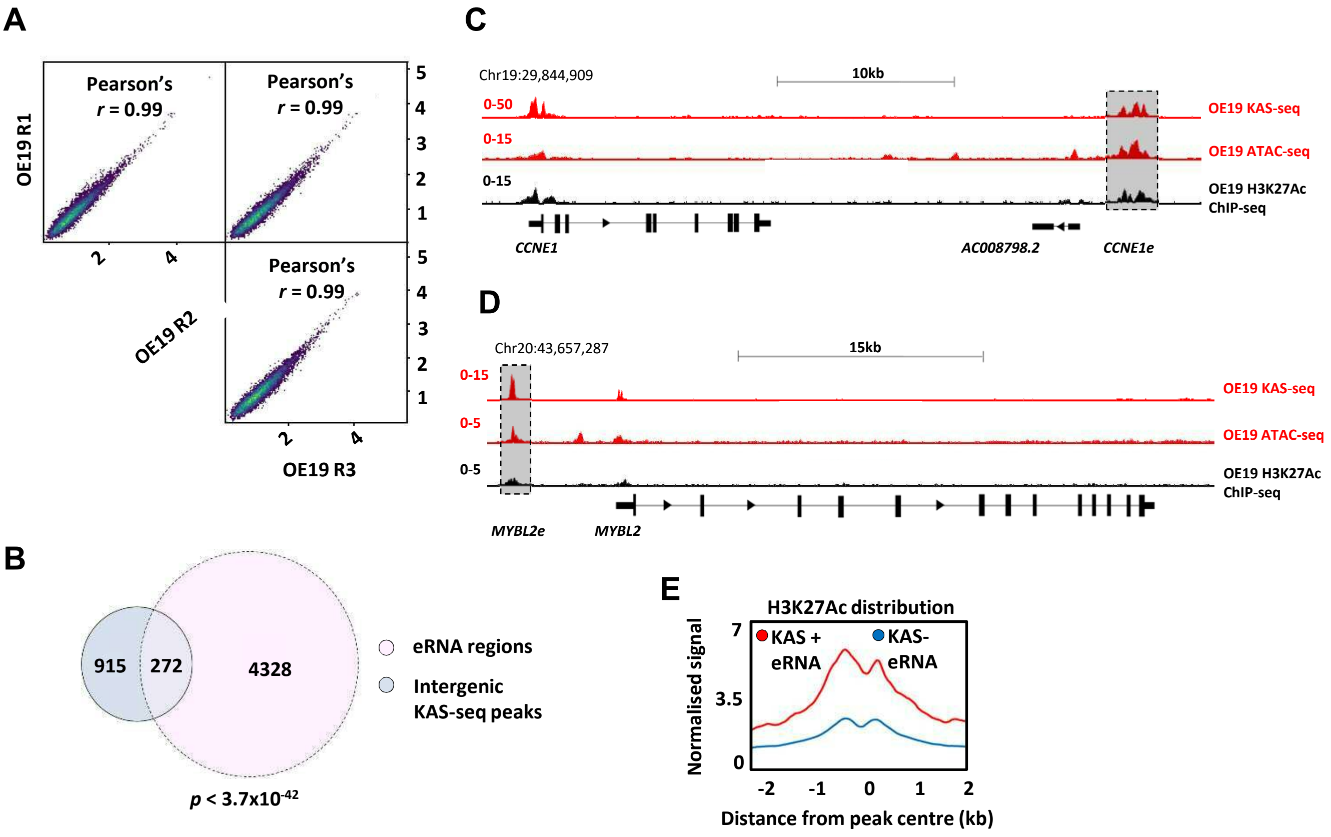


**Figure S3. KAS-seq reveals active transcription at eRNA regions.**
(A) Scatter plots displaying correlation of OE19 KAS-seq biological replicates (Pearson's *r* is shown). (B) Venn-diagram of overlap between 4,600 eRNA regions and intergenic OE19 KAS-seq peaks (*p*-value is shown; hypergeometric test). (C and D) Genome browser views of OE19 KAS- and ATAC-seq data, and OE19 H3K27ac ChIP-seq at the *CCNE1* (C) and *MYBL2* (D) loci with the *CCNE1e* and *MYBL2* eRNAs highlighted. (E) Metaplots of H3K27ac ChIP-seq signal in OE19 cells at 272 KAS+ eRNA regions compared to 4,328 KAS- eRNA regions.

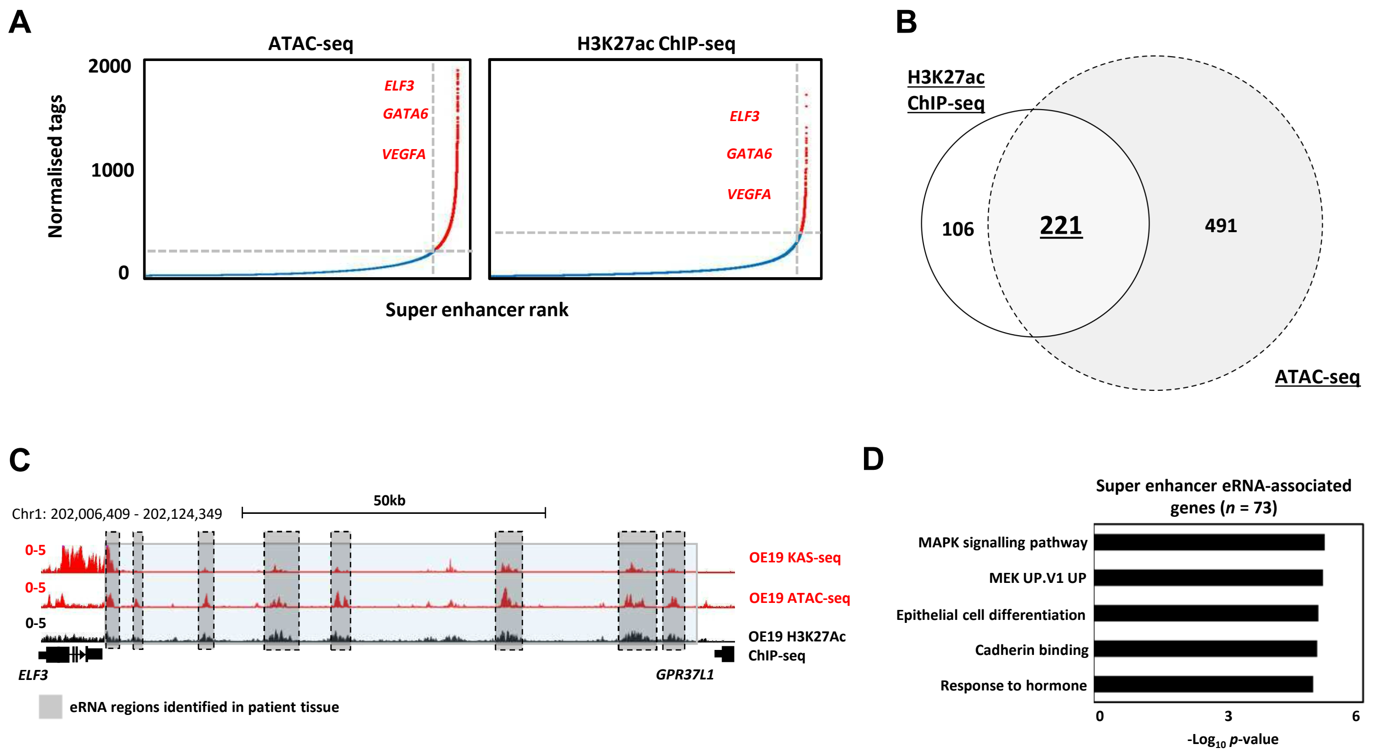


**Figure S4. eRNAs are associated with super enhancers.**
(A) Scatter plots displaying super enhancer stitching on OE19 H3K27Ac ChIP-seq (left) and OE19 ATAC-seq (right) using HOMER. (B) Venn-diagram of overlap between super enhancers identified in OE19 ATAC-seq and OE19 H3K27Ac ChIP-seq (C) Genome browser view of KAS-seq, ATAC-seq data, and H3K27ac ChIP-seq in OE19 cells and eRNAs detected in OAC patient samples at the *ELF3* locus, with the *ELF3* super enhancer highlighted. Putative enhancer regions defined by eRNA expression in OAC patient tissue are shaded. (D) GO-term analysis of genes annotated to high-confidence super enhancers associated with eRNAs.

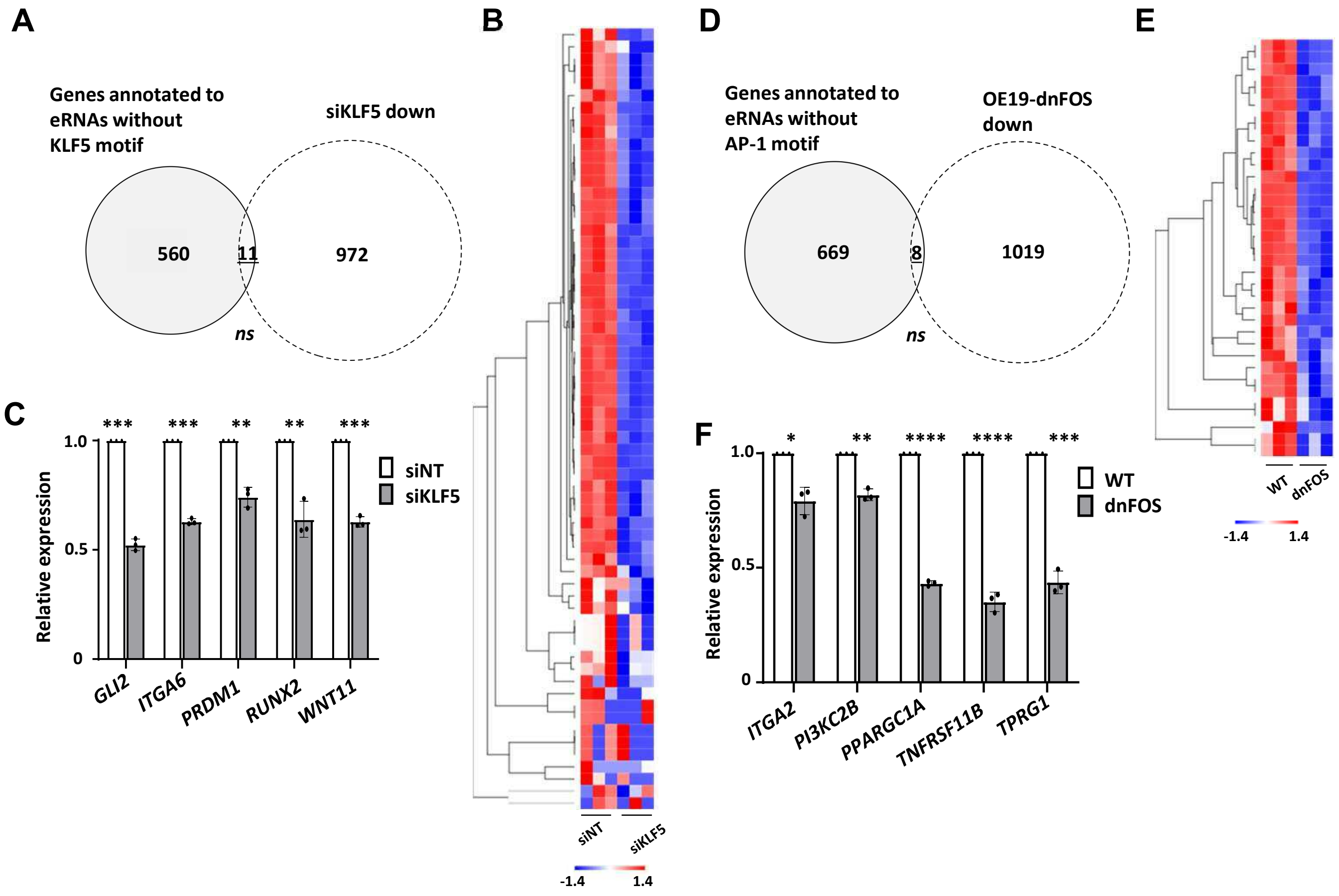


**Figure S5. Association of eRNA regions with potential regulatory transcription factors.** (A) Venn diagram displaying overlap between genes annotated to KLF5 motif lacking eRNAs with genes downregulated upon siKLF5 treatment ($Log_2FC \geq 1.0$, < padj = 0.05) in OE19 cells (*p*-value is non-significant {ns}; Fisher's exact test). (B) Heatmap and hierarchical clustering of siNT (n=3) and siKLF5 (n=3) RNA-seq samples according to row z-score normalised expression of KLF5 eRNA-associated genes which are downregulated upon siKLF5. (C) Bar graphs displaying difference in expression of five KLF5 eRNA-associated genes which are downregulated upon siKLF5 (*** = $p < 0.001$; ** = $p < 0.01$; Welch's t-test). (D) Venn diagram displaying overlap between genes annotated to AP-1 motif lacking eRNAs with genes downregulated upon dnFOS induction ($Log_2FC \geq 0.5$, < padj = 0.05) in OE19 cells (*p*-value is non-significant {ns}; Fisher's exact test). (E) Heatmap and hierarchical clustering of wild-type (n=3) and OE19-dnFOS (n=3) RNA-seq samples according to row z-score normalised expression of AP-1 eRNA-associated genes which are downregulated upon dnFOS. (F) Bar graphs displaying difference in expression of five AP-1 eRNA-associated genes which are downregulated upon dnFOS (**** = $p < 0.0001$; *** = $p < 0.001$; ** = $p < 0.01$; * = $p < 0.05$; Welch's t-test). .

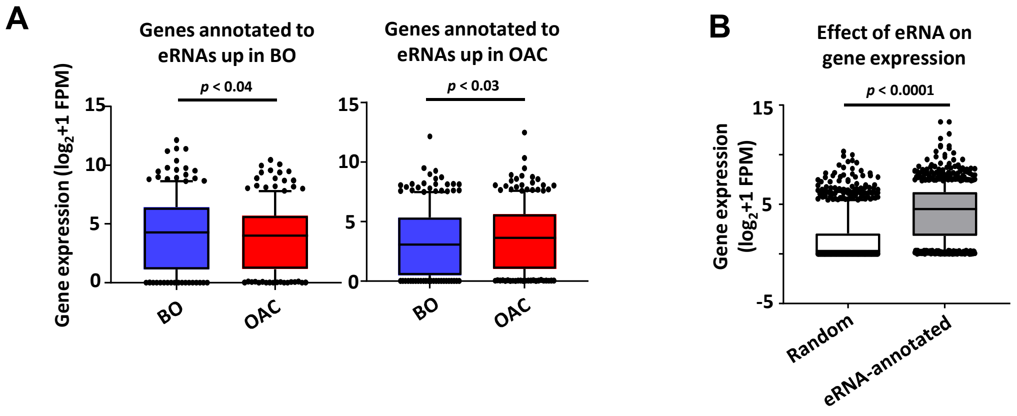


**Figure S6. Association of eRNA regions with potential target genes.** (A) Box plots comparing the expression of genes annotated to eRNAs differentially expressed in BO (left) or OAC (right) in BO and OAC patient tissue total RNA-seq samples from the Maag dataset (Maag et al., 2017) (p-value is shown; Welch's t-test). (B) Box plots comparing gene expression of 1,000 randomly selected genes against genes annotated to eRNAs in the OCCAMs dataset (n = 973; p-value is shown; Welch's t-test).

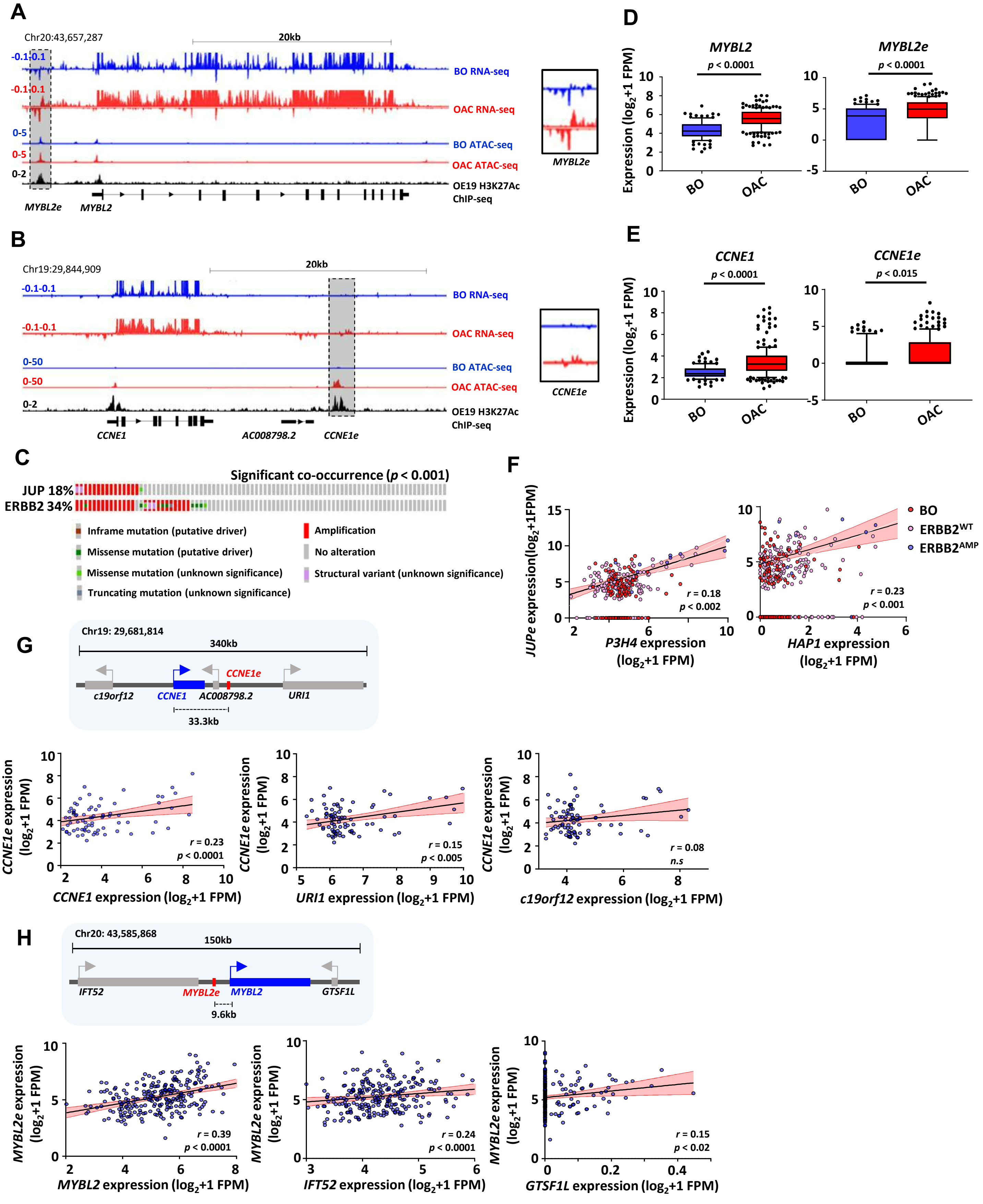


**Figure S7. eRNA regions identify *JUP*, *CCNE1* and *MYBL2* as a candidate target genes.**
(A and B) Genome browser views of BO and OAC patient tissue ATAC- and total RNA-seq data, and H3K27ac ChIP-seq in OE19 cells, at the *MYBL2* (A) and *CCNE1* (B) loci with the *MYBL2e and CCNE1e* eRNAs highlighted and inset. (C) Oncoprint displaying mutational status of *JUP* and *ERBB2* for OAC patients in the TCGA PanCancer Atlas dataset (p-value is shown; one-sided Fisher's exact test). (D and E) Box plots comparing the expression of (D) *MYBL2* (left) and *MYBL2e* (right) or (E) *CCNE1* (left) and *CCNE1e* (right) in BO (n = 108) and OAC (n = 210) patient tissue total RNA-seq samples (p-value is shown; Welch's t-test). (F) Correlation of *JUPe* and (left) *P3H4* or (right) *HAP1* expression across BO (n = 108), $ERBB2^{WT}$ (n = 193) and $ERBB2^{AMP}$ (n = 17) OAC patient tissue total RNA-seq samples (Spearman's r and p-value are shown; Spearman's rank correlation test). (G and H) Correlation of eRNAs and transcripts for (G) *CCNE1e* and *CCNE1* (left) , *URI1* (middle) and *c19orf12* (right) expression and (H) *MYBL2e* and *MYBL2* (left), *IFT52* (middle) and *GTSF1L* (right) expression BO (n = 108) and OAC (n = 210) patient tissue total RNA-seq samples (Spearman's r and p-value are shown; Spearman's rank correlation test). Schematics display the relative locations of putative eRNA region target genes and nearest neighbours (top) .

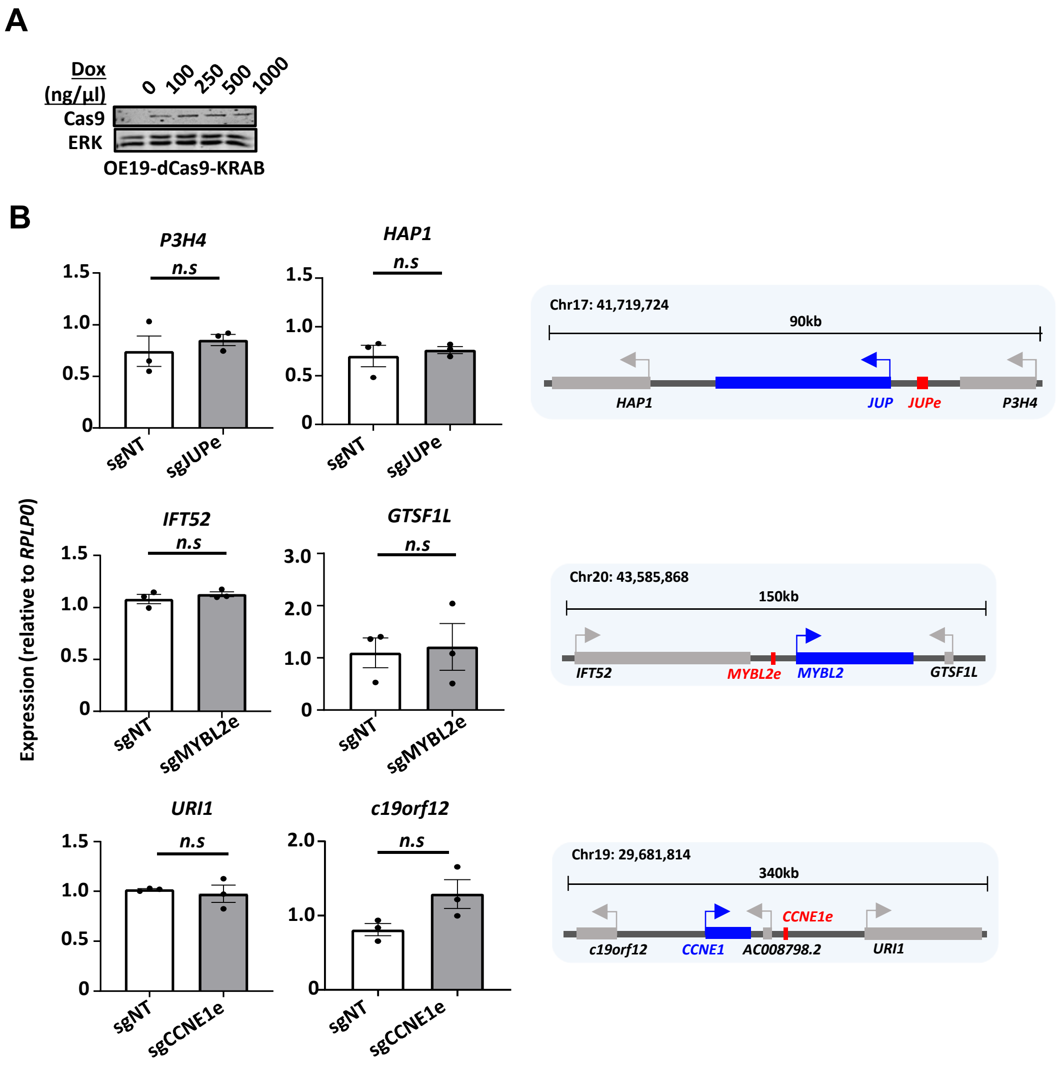


**Figure S8.** ***In vitro*** **interrogation of eRNA regions confirms production and association with cancer-associated processes.**
(A) Western blot showing induction of Cas9 in OE19-dCas9-KRAB cells upon doxycycline treatment. (B) Bar graphs displaying difference in expression of (top) *P3H4* and *HAP1* upon sgJUPe treatment, (middle) *IFT52* and *GTSF1L* upon sgMYBL2e treatment and (bottom) *URI1* and *c19orf12* upon sgCCN1E knockdown, in OE19-dCas9-KRAB cells using RT-qPCR (*p*-value is shown; Welch's t-test).

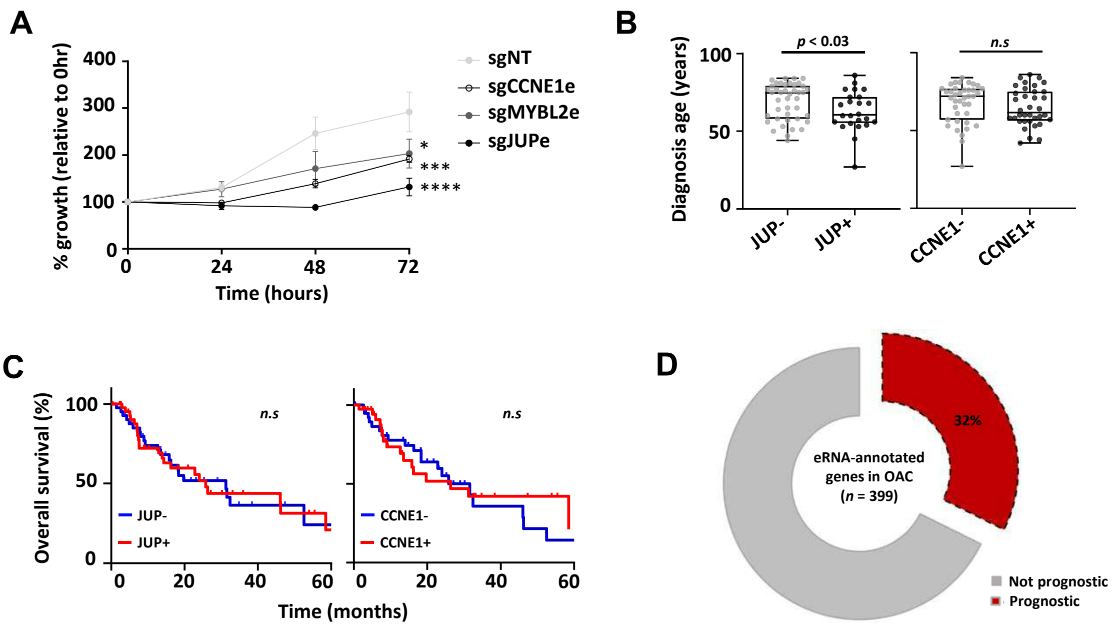


**Figure S9. Biological and clinical relevance of eRNAs and their target genes.**
(A) Growth curves comparing difference in growth in OE19-dCas9-KRAB cells upon indicated sgRNA treatment, assessed by crystal violet assay (* = $p < 0.05$, *** = $p < 0.001$, **** = $p < 0.0001$; two-way ANOVA). (B) Box plots comparing diagnosis age for OAC patients with low and high (left) *JUP* and (right) *CCNE1* in the TCGA PanCancer Atlas dataset (*p*-value is shown; Welch's t-test). (C) Kaplan-Meier plots comparing overall survival between OAC patients with low and high (left) *JUP* expression and (right) *CCNE1* expression in the TCGA PanCancer Atlas dataset (Logrank *p*-value is shown). (D) Number of OAC eRNA-annotated genes that are prognostic for patient survival in the TCGA PanCancer Atlas dataset (Log rank *p*-value <0.05).
